# Rapid screening of engineered microbial therapies in a 3-D multicellular model

**DOI:** 10.1101/491159

**Authors:** Tetsuhiro Harimoto, Zakary S. Singer, Oscar S. Velazquez, Joanna Zhang, Samuel Castro, Taylor E. Hinchliffe, William Mather, Tal Danino

## Abstract

Synthetic biology is transforming therapeutic paradigms by engineering living cells and microbes to intelligently sense and respond to diseases including inflammation^1,2^, infections^3-5^, metabolic disorders^6,7^, and cancer^8,9^. However, the ability to rapidly engineer new therapies far outpaces the throughput of animal-based testing regimes, creating a major bottleneck for clinical translation^10,11^. *In vitro* approaches to address this challenge have been limited in scalability and broad-applicability. Here, we present a bacteria-in-spheroid co-culture (BSCC) platform that simultaneously tests host species, therapeutic payloads and synthetic gene circuits of engineered bacteria within multicellular spheroids over a timescale of weeks. Long-term monitoring of bacterial dynamics and disease progression enables quantitative comparison of critical therapeutic parameters such as efficacy and biocontainment. Specifically, we screen *S. typhimurium* strains expressing and delivering a library of antitumor therapeutic molecules via several synthetic gene circuits. We identify novel candidates exhibiting significant tumor reduction and demonstrate high similarity in their efficacies using a syngeneic mouse model. Lastly, we show that our platform can be expanded to dynamically profile diverse microbial species including *L. monocytogenes, P. mirabilis*, and *E. coli* in various host cell types. This high-throughput framework may serve to accelerate synthetic biology for clinical applications and understanding the host-microbe interactions in disease sites.

## Introduction

The abundance of naturally-occurring microbes living in and on the human body present a myriad of opportunities for engineering microbes to act as *in situ* therapies or diagnostics. As a result, an emerging focus of synthetic biology has been to engineer bacteria to intelligently sense and respond to disease states including inflammation^1,2^, infections^3-5^, metabolic disorders^6,7^, and cancer^8,9^. Notably, many bacteria have been found to selectively colonize tumors *in vivo*, prompting attempts to engineer bacteria as programmable vehicles to deliver anticancer therapeutics^12^. However, a major bottleneck for clinical translation in all these cases has been animal-based testing, which slows iterations of design cycles needed to produce robust microbial systems for *in vivo* applications. Consequently, typical development of engineered bacteria occurs in environments far from *in vivo* conditions, inevitably leading to failure of predicted functions in more stringent native niches^10,13^.

To bridge this gap, *in vitro* platforms have been developed to characterize small-molecule drugs and biologics in more physiologically relevant conditions, with examples including organs-on-a-chip, microfabricated cell patterning, and multicellular spheroids and organoids^14-17^. These systems allow for growth of cells in three-dimensions (3-D), which preserves characteristic features of *in vivo* environments such as gradients in oxygen and metabolites, cell-cell interactions, and intra-cellular variations, as compared to 2-D monolayer cultures^18^. Furthermore, analyses such as drug distribution and spatially heterogeneous responses can be performed that are otherwise unattainable in monolayer systems^19^. For bacterial therapies, three-dimensional disease models could provide an ideal testbed for quantitatively monitoring bacterial localization and circuit dynamics that are critical for accurate estimation of safety and efficacy *in vivo*.

Due to the rapid proliferation rate of bacteria, long-term co-culturing with mammalian cells has been a challenge, limiting assays to short time periods or necessitating the use of heat-killed bacteria^20,21^. Previous attempts to control the imbalance in growth rates have fluidically controlled excess bacteria, directly injected bacteria into multicellular aggregates, or utilized obligate anaerobes to prevent overgrowth^22-24^. As a result, these systems expectedly increase technical complexity and restrict species types, reducing throughput and broad-applicability, respectively. Furthermore, long-term analysis of bacteria circuit dynamics has yet to be employed in multicellular co-cultures or applied towards therapeutic development^21,22^. In order to narrow down the large space of microbial therapy candidates, simple and high-throughput 3-D testing platforms are needed to accelerate development for *in vivo* applications.

Here we present a bacteria-in-spheroid co-culture (BSCC) platform to test a large number of engineered microbial therapies that can be created from combinations of genetic circuits, therapeutic payloads, and host species in 3-D multicellular spheroids (Fig. 1A). By selectively confining bacterial growth within spheroids, we enable parallel and stable co-culture with diverse bacteria and cell types. Using the BSCC system, we screened clinically-relevant *S. Typhimurium* delivering a library of anticancer molecules via synthetic gene circuits and identified novel therapies for *in vivo* applications.

**Fig. 1.**
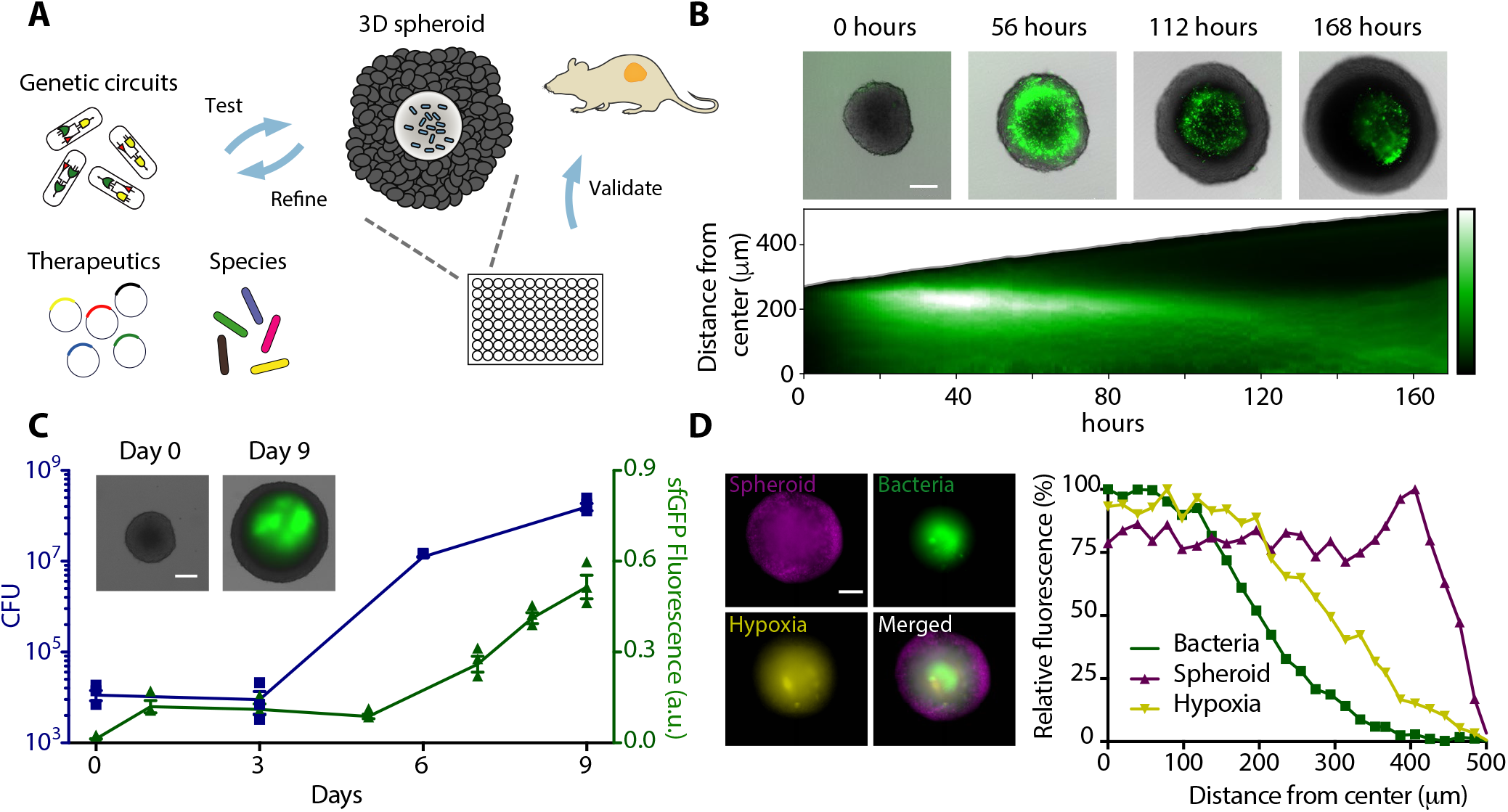
Design and characterization of bacteria-in-spheroid co-culture (BSCC) platform. (A) Schematic of workflow for rapid testing of synthetic gene circuits, therapeutic payloads, and host species using BSCC. Bacteria selectively colonize multicellular spheroids *in vitro* where they are screened for therapeutic efficacy and then are validated in mouse tumor models. (B) *S. typhimurium* and tumor spheroid growth over time. (top) Representative time series images of sfGFP expressing bacteria in CT26 tumor spheroids. Scale bar, 200 μm. (bottom) The corresponding space-time diagram showing radially averaged fluorescence intensity of sfGFP-expressing bacteria. The gray boundary indicates the edge of the spheroid. (C) Total sfGFP of bacteria (green) and colony forming units (CFU, blue) over time. Day 0 values are after inoculation and washing of bacteria. Error bars indicate ± s.e. averaged over three measurements. Scale bar, 200 μm. a.u., arbitrary unit. (inset) Representative images from this data. (D) (left) Spatial distribution of *S. typhimurium* in a spheroid 14 days post colonization. CT26 tumor spheroids (iRFP, magenta) develop hypoxic regions in the spheroid core shown by hypoxic probe dye (yellow). *S. typhimurium* (sfGFP, green) grow inside of the spheroid core. Scale bar, 200 μm. (right) Distribution of fluorescence intensity of tumor cells, bacteria, and hypoxic regions from the center of spheroid.

## Results

### Establishing A Stable Bacteria Co-Culture System in Multicellular Spheroids

To rapidly screen microbial therapies, we generated 3-D multicellular spheroids in 96-well plates for parallel testing^16^ (Movie S1). We first characterized attenuated *S. typhimurium*, a model bacterium well-characterized for its tumor colonization and anticancer applications *in vivo*^12,25^, and constitutively expressed sfGFP to track its dynamics. Following a short incubation of bacteria with spheroids, we screened several antimicrobial agents and found that optimal concentrations of gentamicin confined bacteria within spheroids without causing cell toxicity (Fig. S1 and Table S1). Here, bacteria infiltrated spheroids and subsequently localized and proliferated in the spheroid (Movie S2). To investigate the spatiotemporal dynamics of bacteria in detail, we used automated image analysis to quantify the distribution of fluorescence intensity within the spheroid over time. Bacteria fluorescence was observed to increase near the spheroid boundary (40 hours), reach a steady state level of fluorescence (110 hours), and eventually grow but remain contained within the spheroid (140 hours) (Fig. 1B and Fig. S2). Bacterial proliferation over time was confirmed by plating dissociated spheroids on selective agar (Fig. 1C). Bacteria achieved stable colonization for up to 2 weeks and localized to necrotic and hypoxic regions of spheroids (Fig. 1D and Fig. S2), analogous to bacteria colonization conditions *in vivo*^26^. These results suggest our BSCC platform enables long-term, quantitative characterization of bacterial dynamics within physiologically-relevant 3-D models in a highly parallel manner.

### Quantitatively Monitoring Synthetic Gene Circuit Dynamics in BSCC

We first sought to characterize gene circuit dynamics relevant to *in vivo* therapeutic delivery using the BSCC platform. We constructed an acyl-homoserine lactone (AHL) inducible *luxI* promoter (Fig. 2A) and characterized circuit dynamics by monitoring downstream production of sfGFP from *S. typhimurium* within the spheroid. We applied 10 μM AHL to spheroids after bacteria localized within the spheroid, and observed >400% induction in sfGFP signal that reached steady-state expression within 10 hours (Fig. 2A, Fig. S3 and Movie S3), demonstrating the ability to rapidly trigger therapeutic production within a tumor environment. To sustain therapeutic production without repeated infusion of chemical inducer, we next sought to engineer positive feedback from *luxI* to build a quorum-sensing (QS) circuit (Fig. 2B). This circuit enables self-triggered gene expression when bacteria reach a critical density within the tumor core^27^. After 140 hours of bacterial colonization, we observed a sharp increase in sfGFP production, demonstrating QS activation (Fig. 2B, Fig. S3 and Movie S4). Although sfGFP signal continued to increase over the next 10 hours, maximum induction was approximately 2-fold less than the inducible system (Fig. S3), likely due to reduced levels of local AHL concentration. To enhance drug release from bacteria, we incorporated quorum-mediated bacterial lysis. We modified a recently created synchronized lysis circuit (SLC, Fig. 2C), which produces periodic cycles of self-lysis of host bacteria under QS control to efficiently release therapeutics into the surrounding environment^28^. To minimize bacterial lysis before tumor colonization, we designed the circuit in a single operon on a low copy number plasmid. Upon bacterial colonization of the spheroid, we detected fluctuations in fluorescence from SLC bacteria, indicating periodic lysis (Fig. 2C, Fig. S3 and Movie S5). In addition, spatiotemporal analysis indicated localization of bacteria in the inner spheroid core, while total sfGFP signal did not increase up to 170 hours of bacterial colonization (Fig. 2C and Fig. S4), suggesting a smaller and spatially-restricted bacterial population due to SLC. When gentamicin was removed from the media, we found that SLC maintained containment inside spheroids 2-fold longer than a non-lysing inducible circuit (Fig. S4).

**Fig. 2.**
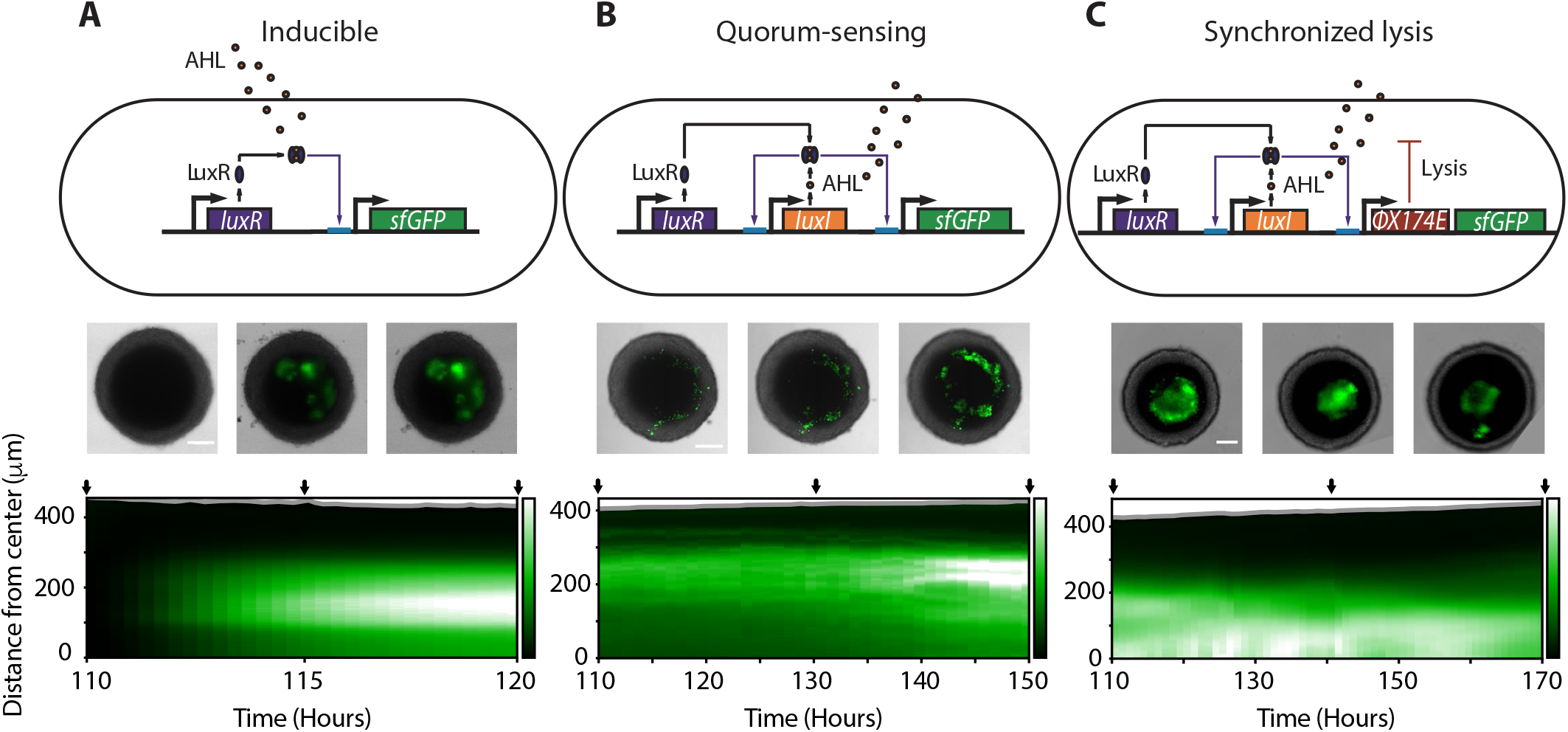
Characterizing synthetic gene circuits using the BSCC platform. (A) Inducible circuit consisting of the *luxI* promoter driving sfGFP activated via externally supplied AHL. (B) Quorum-sensing (QS) circuit contains the *luxI* promoter regulated production of autoinducer AHL, leading to self-activation via positive feedback. (C) Synchronized lysis circuit (SLC) carries an additional lysis gene under the control of *luxI* promoter to achieve periodic lysis. (top) Synthetic gene circuit diagrams. (middle) Representative images of bacteria in tumor spheroids for each gene circuit. Scale bar, 200 μm. (bottom) Space-time diagram showing fluorescence intensity of sfGFP expressing bacteria radially averaged over time in spheroids. Black arrows correspond to images above. *LuxR* is constitutively expressed in all systems.

### Screening Efficacy of Therapeutic Payloads in BSCC

Since the vast majority of bacterial therapy studies test a small number of therapeutic payloads that are rarely compared to one another^12^, we sought to rapidly identify and compare novel bacterial therapeutics using the BSCC platform. We created a library of therapeutics including previously uncharacterized bacterial toxins and anti-cancer peptides (Table S2). Next, we tested bacterial therapy by selectively expressing therapeutics from bacteria within spheroids using the inducible system. After adding AHL to spheroids that contain bacteria, we observed varying degree of reduction in tumor spheroid growth (Fig. 3A). The three candidates that exhibited the highest reduction in spheroid growth were azurin^29^, theta-toxin^30^ and hemolysin E^31^. The former is a pro-apoptotic protein that demonstrated limited efficacy in monolayers. The latter two are pore-forming toxins, of which theta-toxin has not yet been studied as an engineered bacterial therapeutic to our knowledge. Histopathological analysis of treated spheroids revealed higher levels of tumor cell death following treatment with bacteria expressing pore-forming toxins compared to treatment with control bacteria (Fig. S5).

**Fig. 3.**
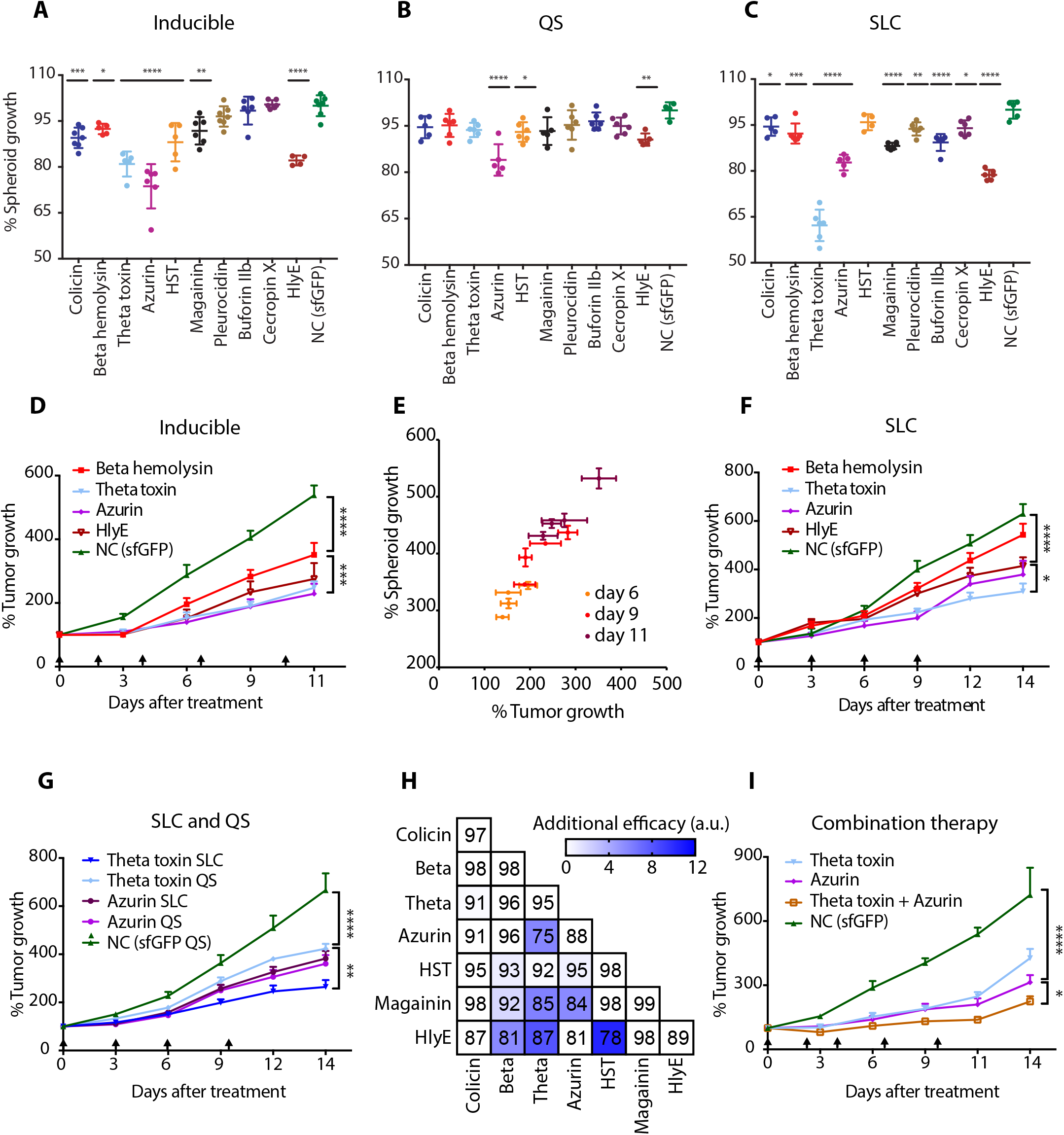
Rapid screening of engineered bacterial therapeutics in BSCC platform and assessment of efficacy *in vivo*. (A-C) Percent spheroid growth 10 days post administration of engineered *S. typhimurium* expressing various anticancer agents with an inducible circuit after (A) adding 10nM AHL post bacteria colonization (****P<0.0001, ***P=0.0002, **P=0.0086, *P=0.0301 compared to control, one-way ANOVA with Bonferroni post-test), (B) quorum-sensing (QS) circuit (****P<0.0001, **P=0.0022, *P=0.0344 compared to control, one-way ANOVA with Bonferroni post-test), and (C) synchronized lysis circuit (SLC) (****P<0.0001, ***P=0.0005, **P=0.0075, *P=0.0394 colicin, *P=0.0015 cecropin X compared to control, oneway ANOVA with Bonferroni post-test). Values are normalized by control (bacteria expressing sfGFP). Error bars indicate ± s.e. averaged over four or more samples. HST, heat stable enterotoxin. HlyE, hemolysin E. NC, negative control. (D) Percent tumor growth over time for subcutaneous CT26 tumor bearing mice injected with *S. typhimurium* carrying inducible gene circuit expressing beta hemolysin, theta-toxin, azurin, hemolysin E, and sfGFP (control) (****P<0.0001, ***P=0.0003, two-way ANOVA with Bonferroni post-test, n=8,7,10,7,5 tumors respectively, error bars show s.e.). (E) Analysis of correlation between tumor growth over time in spheroid and *in vivo* model (Linear regression, R^2^=0.92). (F) Percent tumor growth over time for subcutaneous CT26 tumor bearing mice injected with bacteria carrying SLC expressing beta hemolysin, theta-toxin, azurin, hemolysin E, and sfGFP (control) (****P<0.0001, *P=0.0363, two-way ANOVA with Bonferroni post-test, n=8,9,6,8,8 tumors respectively, error bars show s.e.). (G) Percent tumor growth over time for subcutaneous CT26 tumor bearing mice injected with *S. typhimurium* carrying SLC or QS circuit expressing theta-toxin, azurin, and sfGFP (control) (****P<0.0001, **P=0.0018, two-way ANOVA with Bonferroni post-test, n=8,5,10,10,9 tumors respectively, error bars show s.e.). Black arrows indicate bacteria injections in mice. (H) Efficacy of combinatorial therapy in BSCC platform. The numbers in each box indicate average percent spheroid growth over two or more measurements 8 days post bacteria administration. Colors indicate additional efficacy of combinatorial therapy. Additional efficacy represents increased efficacy compared to additive effect alone (see Methods). (I) Percent tumor growth over time for subcutaneous CT26 tumor bearing mice injected with *S. typhimurium* carrying inducible circuit expressing theta-toxin, azurin, combination (theta-toxin and azurin) and sfGFP (control) (****P<0.0001, *P=0.0117, two-way ANOVA with Bonferroni post-test, n=7,9,9,5 tumors respectively, error bars show s.e.).

To compare the effects of genetic circuit on efficacy, we tested expression of the therapeutic library across inducible, QS, and SLC systems in the BSCC platform. Incorporating the QS circuit into bacteria expressing the therapeutic library produced limited efficacy in tumor spheroids (Fig. 3B), corroborating with our observation that quorum activation is limited compared to inducible circuits. In contrast, many therapeutics displayed significant efficacy when expressed from the SLC system (Fig. 3C). Comparing across the therapeutic library, we identified theta-toxin as a potent therapy when combined with SLC, resulting approximately 40% reduction in tumor spheroid growth (Fig. 3C). Given the toxicity of theta-toxin to host bacteria (Fig. S6), we varied AHL in the inducible circuit from 10 nM to 0 nM and found that decreasing inducer concentration did not increase efficacy (Fig. S6). Therefore, we reasoned that theta-toxin expression from SLC exhibited higher therapeutic efficacy compared to the inducible circuit due to efficient release.

### Validating Efficacy of Bacterial Therapies in an Animal Model

To determine the predictive ability of the BSCC system, we assessed engineered bacteria in a syngeneic, hind-flank tumor mouse model harboring CT-26 cells identical to those used to generate spheroids. We investigated inducible therapeutics with predicted high efficacy (azurin, theta-toxin and hemolysin E), moderate efficacy (beta-hemolysin), and control (sfGFP). Approximately 3-fold reduced tumor growth was observed by bacterial therapeutics identified as highly effective from the *in vitro* screen compared to the control treatment (Fig. 3D). Tracking therapeutic responses over time revealed a high degree of similarity in trends of efficacy between BSCC and *in vivo* results (Fig. 3E). In contrast, results from bacteria co-cultured with a monolayer of the same CT-26 cells failed to predict trends of efficacy *in vivo* (Fig. S7). We performed histopathological analysis on treated tumors at trial termination. Consistent with the spheroid results, tumors treated with theta-toxin and hemolysin E showed increased cell death relative to tumors treated with control bacteria (Fig. S8).

We next examined the QS and SLC circuits, and combination therapy in mouse tumor models. We found that efficacy of therapeutics expressed under SLC control *in vivo* corresponded to those from the spheroid screen (Fig. 3F). Comparing efficacy of SLC and QS circuits, theta-toxin from SLC exerted the strongest response, similar to the results from the BSCC platform (Fig. 3G). Furthermore, overall animal health as measured by weight drop improved with SLC compared to bacteria carrying inducible circuits, similar to the observation of enhanced biocontainment by SLC in the BSCC platform (Fig. S9). Leveraging the high-throughput capability of the BSCC, we also investigated combination therapy. Applying pairwise combinations of bacteria carrying inducible circuits at equal proportions in the BSCC, we found theta-toxin and azurin produced the most effective combination (Fig. 3H), exhibiting higher therapeutic efficacy than the additive effect of each individual therapy. This combination therapy *in vivo* exerted significantly stronger anti-tumor effects than either therapeutic alone, yielding a 4-fold reduction in tumor graft growth compared to control (Fig. 3I). These findings indicate the BSCC platform predictively identifies potent genetic circuits and therapeutic combinations in a highly parallel manner.

### Expanding Applicability of BSCC to Diverse Bacterial Strains, Species and Cell Types

Given existing 3D co-culture models for bacterial therapy are typically limited to particular types of bacteria and cells^24,32^, we assessed the ability of the BSCC platform to co-culture diverse bacterial strains, species and host tissue types. First, we examined the colonization capacity and sfGFP expression level of clinically-relevant strains of *S. typhimurium*: ELH430 (SL1344 *phoPQ*-), SL7207 (SL1344 *aroA*-), ELH1301 (SL1344*phoPQ-/aroA*-)^33,34^, and VNP20009 (14028S *msbB-/purI-*), which has been used in clinical trials^25^. All strains successfully colonized tumor spheroids, demonstrating stable co-culture (Fig. S10). ELH1301 exhibited the highest colonization and sfGFP expression levels within spheroids (Fig. S10). Analysis of dynamics revealed increased initial invasion by ELH1301 compared to others (Fig. 4A), possibly contributing to its high colonization capacity. Next, we investigated colonization of *L. monocytogenes, P. mirabilis*, and *E. coli*, bacterial species previously tested for cancer therapy^26,35^. Optimizing bacterial inoculation density, incubation time, and gentamicin concentrations (Table S3), we established long-term growth of all bacterial species, indicated by increase in sfGFP fluorescence and CFU from spheroids over multiple days (Fig. 4B-D). Spatiotemporal analysis allowed for comparison of fluorescence distribution within spheroids and respective dynamics. *L. monocytogenes* displayed a notably wide area of tumor colonization (Fig. 4B), possibly because of its cell-to-cell spreading ability within tumor mass^36^. *P. mirabilis* reached the spheroid core at earlier time compared to other bacteria (Fig. 4C), which might be attributable to its swarming ability. *E. coli* displayed a sharp increase in number at day 6 within the spheroid core (Fig. 4D), demonstrating similar dynamics to *S. typhimurium*. In addition to the ability to test host species, the BSCC platform can be used to study different tissue types. To do this, we generated spheroids derived from both human and mouse colorectal (HT-29 and CT-26/MC-26) and breast (MCF-7 and 4T1) cells. We observed increasing sfGFP fluorescence from *S. typhimurium* over time (Fig. S11), demonstrating growth of bacteria in various cell lines. As efforts to integrate additional bacterial species for therapeutics continue to expand, we anticipate that the modularity of the BSCC system allows rapid evaluation of broad microbial therapies and disease modalities.

**Fig. 4.**
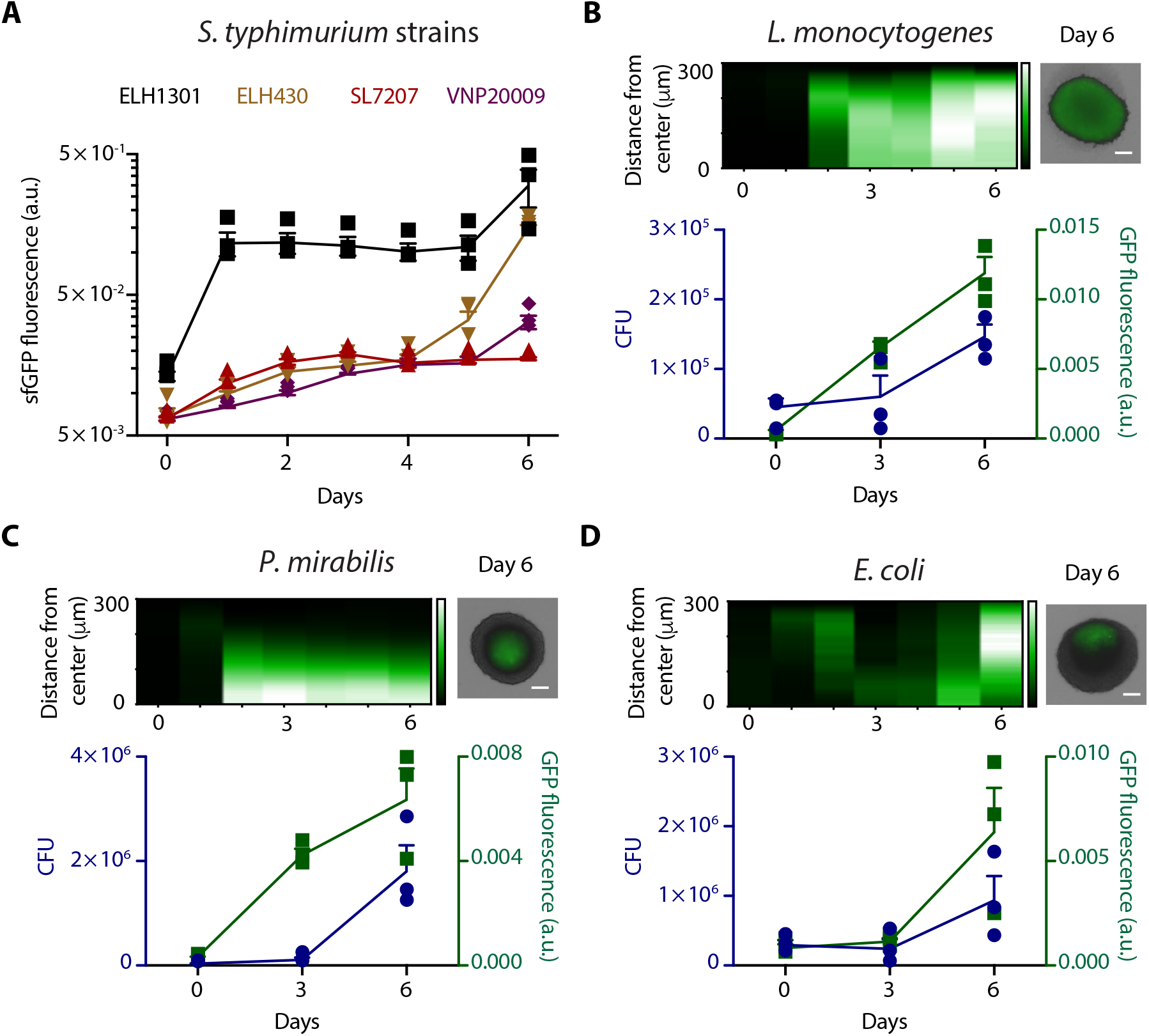
Co-culture of diverse bacterial strains and species in BSCC platform. (A) sfGFP fluorescence of *S. typhimurium* strains within spheroids over time. (B-D) Colonization dynamics of (B) *L. monocytogenes*, (C) *P. mirabilis*, (D) *E. coli* within spheroids. (left) Space-time diagram showing fluorescence intensity of sfGFP expressing bacteria radially averaged over time. (right) Typical images of sfGFP expressing bacteria in spheroids 6 days post bacteria administration. Scale bar, 100 μm. (bottom) Total sfGFP of bacteria (green) colony forming units (CFU, blue) over time. Day 0 values are after inoculation and washing of bacteria. Scale bar, 100 μm. a.u., arbitrary unit. Error bars indicate ± s.e.

## Discussion

We present an approach to simultaneously profile a vast number of engineered bacteria in a physiologically-relevant, 3-D multicellular system. By utilizing a scalable methodology, the BSCC platform enables rapid ‘build-test’ cycles of bacterial dynamics, therapeutics, and species for *in vivo* applications of synthetic biology. The key features of our system are: 1) quantification of bacterial population and circuit dynamics in a 3-D disease model, 2) accurate prediction of long-term disease progression *in vivo* from *in vitro* screening, and 3) a simple and broadly-applicable system for high-throughput development of novel therapies.

The BSSCC platform has several advantages over existing co-culture of microbes and mammalian cells, which have been of limited utility for the development of bacterial therapeutics. Traditional co-culture with monolayer of cells typically rely on frequent dilution or physical compartmentalization to balance the rapid growth of bacteria^21^, and often do not mimic the stoichiometry, geometry, or environmental conditions observed *in vivo*. More complex *in vitro* platforms including spheroids and organoids have been introduced to recapitulate the 3-D spatial orientation seen *in vivo*, and have been used as a high-throughput screening tool for molecular-based therapeutics^16,37^. However, these systems have yet to be developed for testing bacterial therapies. The BSCC system combines the advantages of 3-D multicellular co-cultures with high-throughput drug screening and allows for a simple and broadly-applicable system to characterize bacterial therapies.

One of the promising applications of the BSCC technology is the ability to rapidly screen large libraries of therapeutics expressed by bacteria circuit variants to narrow candidate selection for further development. While previous studies have tested up to 5 bacteria-produced therapeutics in monolayers using bacterial lysate and culture supernatants^38^, our technology enabled screening of ~40 bacterial therapy candidates for efficacy while simultaneously monitoring bacteria dynamics and disease progression over a time scale of weeks. The long-term monitoring aspect is critical to the success of bacterial therapies, as it is necessary to characterize the effects of dynamic therapeutic expression on bacteria and cellular growth in a spatially-dependent manner. In this study, we were able to identify theta-toxin driven by the SLC as an efficacious bacterial therapy candidate compared to several previously-tested toxins. Although further studies are required to explore specific mechanisms involved, our results suggest this may be due to effective therapeutic release and minimization on host toxicity provided by the SLC. Furthermore, the BSCC system identified that theta-toxin combined with azurin was an efficacious combination, providing additional efficacy compared to either single therapy. In future studies, the platform can be used to screen even larger libraries and combinations of 3 to 4 agents as often utilized in chemotherapy regimens^39^. Since multiple bacteria strains expressing different therapeutics will compete for resources within a tumor, reducing the effective dosage of each therapeutic, it has been unclear how combination therapy will perform when increasing the number of bacterial therapies. The BSCC system is precisely suited to quantitatively explore this strategy and determine how specific selection of orthogonally-acting therapeutics can be used to increase efficacy.

While we focused on cancer therapy in this study, the BSCC may be expanded to characterize bacteria-based therapeutics for various diseases, and explore fundamental biological questions about bacteria in host tissue types. For example, recent findings have demonstrated the presence of bacteria in patients with pancreatic tumors, and identified their potential role in degradation of chemotherapies^40^. The BSCC system would be able to provide a high-throughput way to assess the interactions between bacteria, cancer cells, and chemotherapies for further exploration. Additionally, several studies have reported a role for bacteria to augment immunotherapies^41,42^. By incorporating immune cells into the co-culture platform, the BSCC system may also be amenable to investigate bacteria-based immunotherapeutics. Furthermore, while multicellular spheroid models enable a reproducible and high-throughput assay format, other 3-D disease models such as organoids incorporate cells with different lineages that may enable further investigation of host-microbe interactions and efficacy. As the field of engineered cell and microbial therapies evolves with the use of more complex synthetic gene circuits^43,44^ and incorporation of multi-cellular ecosystems^45^, we envision the BSCC platform to accelerate therapeutic development for various diseases towards clinical translation.

## Supporting information

## Acknowledgements

We thank the Herbert Irving Comprehensive Cancer Center Molecular Pathology Shared Resources Facility for help with histological sample processing. We thank N. Arpaia, M. Omar Din, and J. Hasty for their helpful comments and suggestions. This work was supported in part by the Honjo International Foundation Scholarship (T.H.), NIH Ruth L. Kirschstein National Research Service Award (Z.S.), NIH Pathway to Independence Award (R00CA197649-02), DoD Idea Development Award (LC160314), and DoD Era of Hope Scholar Award (BC160541). T.H., Z.S. and T.D. have filed a provisional patent application with the US Patent and Trademark Office related to this work.

## Author contributions

T.H., Z.S., and T.D. conceived and designed the BSCC platform. T.H., Z.S. and O.S.V. performed experiments and established the BSCC platform. T.H. built and tested bacterial strains, gene circuits, and therapeutic library. T.H., O.S.V., J.Z., S.C. and T.E.H. designed and performed *in vivo* experiments. T.H., W.M, and T.D. performed image and data analysis. T.H. and T.D. wrote the manuscript with input from all of the other authors.

## Materials and Methods

### Host strains and culturing

ELH1301 and ELH430 were kindly provided by Dr. Elizabeth Hohmann. SL7207 was kindly provided by Dr. Siegfried Weiss. *L. monocytogenes* was kindly provided by Dr. Eric Pamer. VNP20009, *P. mirabilis* and *E. coli* were obtained from ATCC (202165, 29906, 23506). For full strain information, please refer to the Table S4. *S. typhimurium, P. mirabilis* and *E. coli* were cultured in LB media (Sigma-Aldrich). *L. monocytogenes* was cultured in Brain Heart Infusion media (Fisher Scientific). All bacteria were grown with appropriate antibiotics selection (100 μg ml^-1^ ampicillin, 50 μg ml^-1^ kanamycin, 25 μg ml^-1^ chloramphenicol) at 37 °C. Synchronized lysis circuit strains were cultured with 0.2% glucose for less than 16 hours. The glucose was added in order to decrease expression from the *luxI* promoters.

### Mammalian cells and spheroid generation

The MC-26 cell line was provided by K. Tanabe and B. Fuchs (Massachusetts General Hospital). Mammalian cells were cultured in DMEM/F-12 media with GlutaMAX supplement (Gibco; for HT-29 and MCF-7) or RPMI 1640 media (Gibco; for CT-26, CT-26-iRFP, MC-26, 4T1) and supplemented with 10% fetal bovine serum (Gibco) and 1% penicillin/streptomycin (CellGro), placed inside a tissue culture incubator at 37 °C maintained at 5% CO2. For full cell line information, please refer to the Table S4. CT26 cells were transfected with genomic integration of *iRFP* gene with nuclear localization signal sequence to construct CT26-iRFP cell line. Specifically, pNLS-iRFP670 plasmid was obtained from Addgene (Plasmid #45466). The plasmid was prepared by miniprep of an overnight culture and subsequently transfected into CT26 cells using lipofectamine (Invitrogen). Monoclonal cells were selected by serial limiting dilution into 96-well plates (Falcon). Selected *iRFP*-expressing monoclones were expanded and used to seed spheroids in all corresponding assays.

Tumor spheroids were generated by seeding cells in round-bottom ultra-low attachment 96 well plate (Corning). Each well contains 2,500 CT-26 cells in 100 μl of appropriate media without antibiotics. Number of cells seeded was adjusted for each cell line to maintain a similar diameter of spheroids generated across cell lines (MC-26 2,500 cells, 4T1 5,250 cells, HT-29 5,000 cells, MCF-7 12,500 cells). The plate was centrifuged at 3,000 rcf for 5 minutes to aggregate cells at the bottom of the plate and placed inside a tissue culture incubator for 4 days before co-cultured with bacteria.

### Plasmids and therapeutic library constructions

Plasmids were constructed using Gibson Assembly or using standard restriction digest and ligation cloning and transformed into Mach1 competent cells (Invitrogen). Previously constructed pTD103 sfGFP plasmid containing kanamycin resistance cassette and ColE1 origin of replication was used to characterize the inducible gene circuit^1^. The QS circuit (pTH02) was constructed by switching antibiotic resistance cassette and origin of replication of previously used pTD103 LuxI sfGFP plasmid^2^ to ampicillin and p15A using the modular pZ plasmids^3^. The synchronized lysis circuit (SLC) plasmid (pTH03) was constructed by first amplifying a region containing the constitutively expressed *luxR* gene and *luxI* gene under the control of *luxI* promoter from pTD103 LuxI sfGFP plasmid. Next, the bacteriophage *ϕ* X174E was synthesized from IDT and cloned next to the *luxI* gene with intergenic RBS sequence between genes to allow for operon expression (GAGGCAGATCAA). Finally, the antibiotic resistance cassette and origin of replication was replaced with ampicillin and sc101* from a modular pZ plasmid. Maps of main plasmids used in this study (Fig. S12) are provided.

The therapeutics library was constructed by synthesizing therapeutic genes from IDT, except for the hemolysin E gene obtained via PCR from a plasmid in previous work^4^. Therapeutics were cloned under the control of the *luxI* promoter by replacing *sfGFP* gene in a previously used ColE1 pTD103 sfGFP plasmid to construct therapeutics with inducible control (pTH05) ^1^. To combine QS circuit and SLC with therapeutic expression, plasmids containing these circuits were co-transformed into bacteria and plated on full antibiotics. To make therapeutic characterization comparable between circuits, the QS circuit was swapped to a sc101* origin of replication (pTH06). A detailed table of therapeutics in the library (Table S2 and S4) and main plasmids used in this study (Fig. S12) are provided.

### Bacteria co-culture with tumor monolayer cells

Cell viability experiment was performed in 96-well tissue culture plates (Falcon). CT26 cells were allowed to adhere to the wells for 24 hours before the addition of bacteria. 1 x 10^5^ bacteria carrying therapeutic payloads were co-cultured for 4 hours before media was replaced by new cell culture media containing 50 μg/ml gentamicin (Gibco). After 48 hours, viability was assessed using an MTT assay by measuring the colorimetric output in a TECAN Infinite M200 Pro plate reader.

### Bacteria co-culture with tumor spheroids

Bacteria were cultured in a 37 °C shaker overnight to reach stationary phase before use. 10^6^ CFU *S. typhimurium* were inoculated into wells containing 4-day old tumor spheroids and placed back into the tissue culture incubator. After 2 hours of bacteria inoculation, media was removed and tumor spheroids were washed with 200 μl of PBS repeatedly while leaving spheroids at the bottom of plate. After washing, 200 μl of media containing 2.5 μg/ml gentamicin (Gibco) was added and tumor spheroids were monitored for growth. For long-term experiments, media was replaced every 3-4 days. Schematic of the overall protocol is provided (Fig. S1). Bacterial therapeutics expression was induced at day4 by replacing media containing 10 nM N-(β-Ketocaproyl)-L-homoserine lactone (AHL) (Sigma Aldrich). For other bacteria, identical procedures were followed with modifications in bacterial inoculation density, incubation time, and gentamicin concentration. For full protocol, please refer to Table S3. ELH1301 (SL1344 *phoPQ-/aroA-*) was used for all co-culture experiments unless otherwise noted.

### Bacterial colonization quantification via colony counts

Spheroids containing bacteria were washed with 200 μl of PBS repeatedly while leaving spheroids at the bottom of plate. After washing, spheroids were re-suspended in 100 μl of PBS and homogenized using mechanical dissociation with sterile tips and repeated pipetting. Destruction of spheroids were confirmed by microscopy. Serial 10-fold dilutions of the samples were inoculated on appropriate agar plates.

### Microscopy

Acquisition of spheroid still images was performed with EVOS FL Auto 2 Cell Imaging Systems. The scope and accessories were programmed using the Celleste Imaging Analysis software. For analysis of synthetic gene circuit dynamics, Nikon TiE microscope equipped with Okolab stage top incubator was used to maintain the culture at 37 °C with 5% CO2 for time-lapse movies. The scope and accessories were programmed using the Nikon Elements software and images were taken every 60 minutes. For the acquisition of images, we used an Andor Zyla sCMOS camera. The microscope and acquisition was controlled by the Nikon Elements software. Phase-contrast images were taken at 10x magnification at 50-200ms exposure times. Fluorescent imaging at 10x was performed at 70ms for sfGFP, 30% setting on the Lumencor Spectra-X Light Engine. Further information on the analysis of these images is presented in the Image Analysis section below.

### Image alignment and localized fluorescence measurement for spheroids

Time-course imaging of tumor spheroids required image registration to properly align the resulting images, since the spheroid would both rotate and translate significantly within the field of view. For image registration, we leveraged the popular registration method ORB^5^ in the Python implementation of OpenCV to find matching key points in images at adjacent time points. An implementation of RANSAC^6^ in scikit-image^7^ determined the corresponding Euclidean transform between adjacent images. For each time point, we averaged edge-filtered transmitted light (TL) and edge-filtered sfGFP images to form the input for our registration pipeline, which allowed both TL and sfGFP to inform alignment. Once this method was used to align a time-course experiment, Fiji^8^ was used for data analysis. sfGFP fluorescence trajectories for a spheroid (Fig. S3a) are based on averaged signal of this aligned image set within several circular regions of interest (ROI’s) of the same size. These circular ROI’s are fixed in position and chosen to highlight representative dynamics of the spheroid. Images showing ROI’s used in this study are provided in Fig. S3.

### sfGFP average fluorescence and radial histograms for spheroids

To measure the spatiotemporal dynamics of bacteria invading tumor spheroids, we first found a threshold brightness value for each TL image to distinguish the dark spheroid from the light background. We used scikit-image implementations of two popular thresholding methods: the minimum method^9^ for images taken daily, and Yen’s method^10^ for other images. We then identified the largest region within the resulting threshold-based image mask as the tumor spheroid, and we determined mean intensity of sfGFP fluorescence within this region. To compute radial histograms, we computed mean sfGFP fluorescence for many thin annuli with variable mean radius and centered on the centroid of the spheroid mask region. To compute radial fluorescence in Fig. 1d, fluorescence was calculated based on a line across the radius of the spheroid.

### Animal models

All animal experiments were approved by the Institutional Animal Care and Use Committee (Columbia University, protocol AC-AAAN8002). The protocol requires animals to be euthanized when tumor burden reaches 2 cm in diameter, or under veterinary staff recommendation. Mice were blindly randomized into various groups. Animal experiments were performed on 4-6 weeks-old female BALB/c mice (Taconic Biosciences) with bilateral subcutaneous hind flank tumors from CT26 colorectal cells. The concentration for implantation of the tumor cells was 5x10^7^ cells per ml in RPMI (no phenol red). Cells were injected at a volume of 100 μl per flank, with each implant consisting of 5 x 10^6^ cells. Tumors were grown to an average of approximately 150 mm^3^ before experiments. Tumor volume was quantified using calipers to measure the length, width, and height of each tumor (V = L × W × H). Volumes were normalized to pre-injection values to calculate relative or % tumor growth on a per mouse basis.

### Bacterial administration for *in vivo* experiments

Bacterial strains were grown overnight in LB media containing appropriate antibiotics and 0.2% glucose. A 1:100 dilution into media with antibiotics was started the day of injection and grown until an OD of approximately 0.1. Bacteria were spun down and washed 3 times with sterile PBS before injection into mice. Intratumoral injections of bacteria were performed at a concentration of 5 × 10^8^ cells per ml in PBS with a total volume of 20–40 μl injected per tumor. For bacteria carrying the inducible circuit, 0.5 mL of 10 μM AHL was injected subcutaneously the day after bacterial treatment to induce therapeutic expression. For combination therapy, bacteria cultures were combined after washing with PBS at 1:1 ratio to reach total concentration of 10^9^ cells per ml and injected the total volume of 20-40 μl per tumor.

### Tissue histological analysis

Tumors were extracted from mice at the termination of trials according to protocol. Immediately after extraction, tumor tissues were rinsed with PBS and fixed in 4% paraformaldehyde for 24 hours in 4 °C. Tumor spheroids were fixed for 20 minutes to prevent over fixation. After fixation, the tissues were rinsed with PBS and preserved in 70% ethanol in 4 °C. For histological analysis, tissues were paraffin embedded and sectioned into 5 μl onto slides. Gram staining was performed to confirm presence of bacteria inside the tumors. TUNEL staining was performed to obtain measurement of apoptosis. Tumor cell death was quantified using Fiji software to measure the area of the viable and dead area by setting a pixel threshold to make binary images.

### Statistical analysis

Statistical tests were calculated either in GraphPad Prism 7.0 (Student’s *t*-test and ANOVA) or Microsoft Excel. The details of the statistical tests carried out are indicated in the respective figure legends. Where data were approximately normally distributed, values were compared using either a Student’s *t*-test or one-way ANOVA for single variable, or a twoway ANOVA for two variables with Bonferroni correction for multiple comparisons. Mice were randomized in different groups before experiments.

### Calculation of additional efficacy

We calculated additional efficacy, which represents the increased efficacy observed compared to expected from additive effect alone, based on past calculations of synergy and combination therapies for other drugs^11^. Unique to bacterial therapy combinations, when two different bacteria share the tumor space, each can only grow to half of the possible maximum bacterial volume of a single therapy. Efficacy of a single therapy (for example, therapy A) was calculated by measuring the percent reduction of spheroid growth as a result of treatment with that therapy alone. For example, 80% relative tumor growth after treatment with A was counted as A = 20% efficacy. The expected additive efficacy between two therapies was calculated as (A+B)/2. Thus, to calculate additional efficacy, we subtracted the expected additive efficacy from the measured efficacy from combinatorial treatment with A and B (“AB” term in the equation), with a formula of additional efficacy = AB - (A+B)/2, so that any result greater than 1 indicated the combinatorial therapeutic strategy had greater efficacy than additive effect of the component therapies.

## Supplementary Figures

**Fig. S1.**
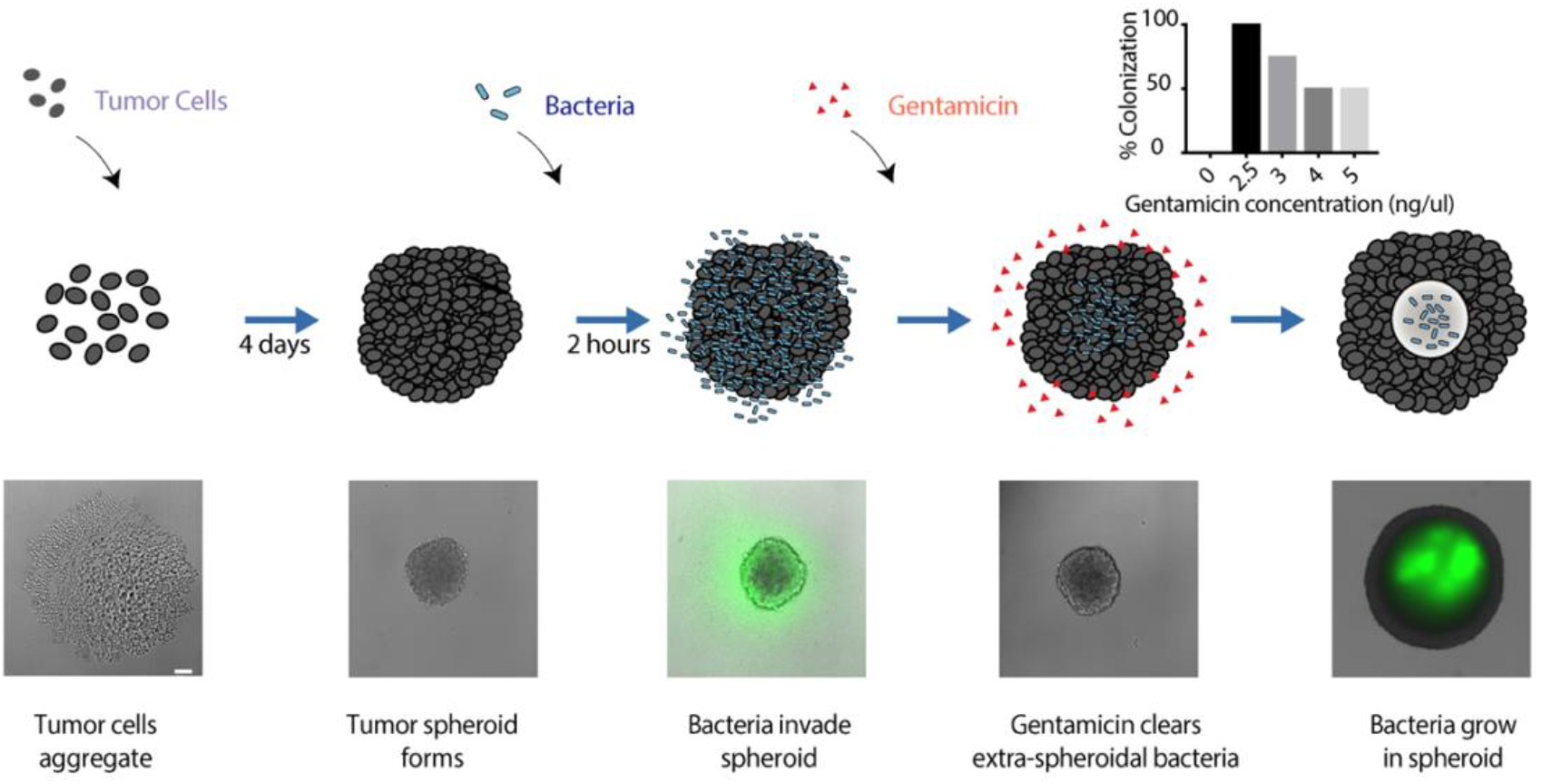
Protocol for establishing *S. typhimurium* spheroid co-cultures. Shown below the schematics are typical images of bacteria (sfGFP, green). Scale bar, 200 μm. (Inset) Screening of gentamicin concentrations for percentage colonization. Colonization probability is calculated out of 10 samples. Detailed co-culture protocol for other species are shown in Table S3.

**Fig. S2.**
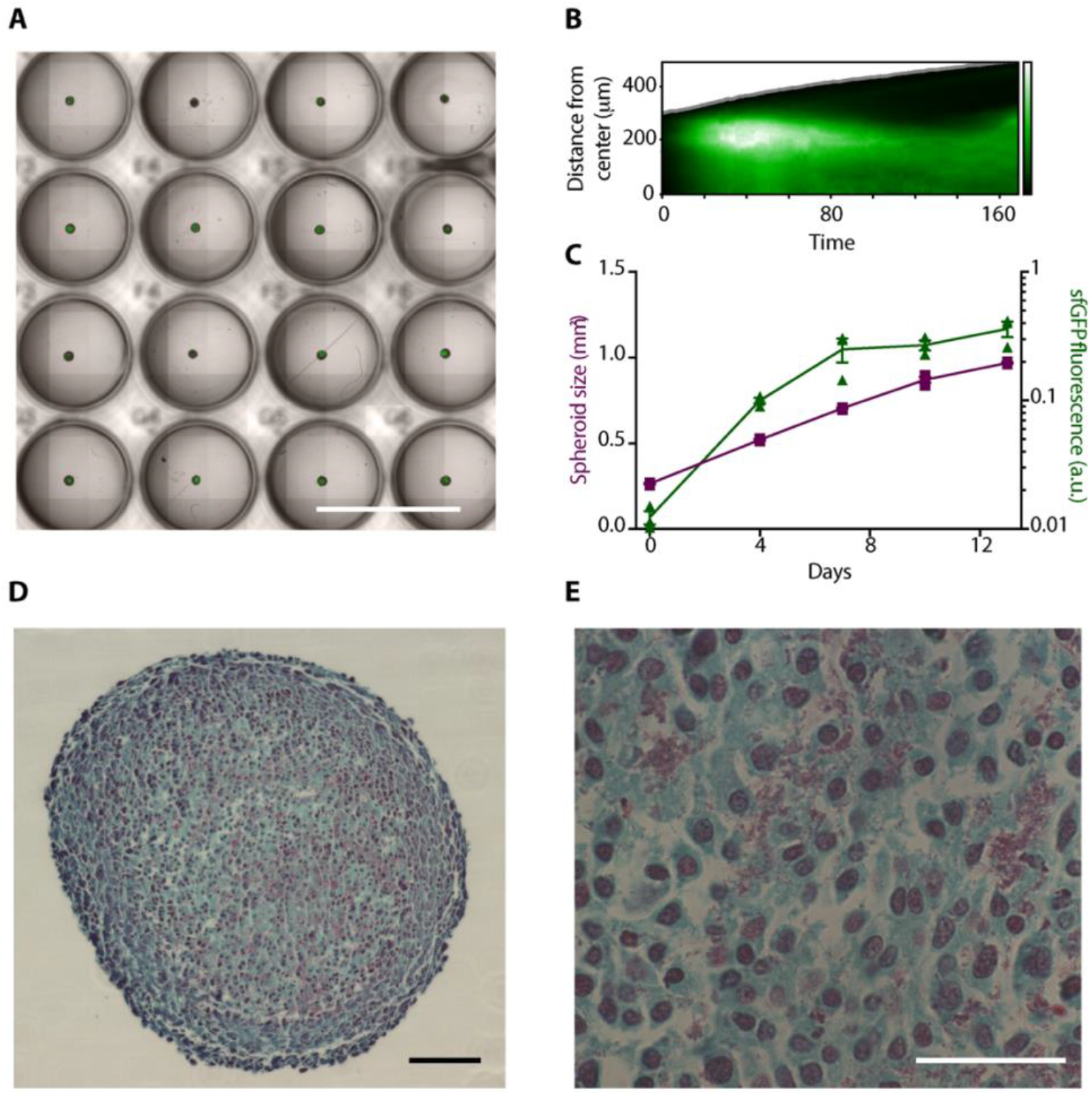
*S. typhimurium* colonization of tumor spheroids. (A) High-throughput co-culture of bacteria with tumor spheroids in 96 well plates. Scale bar, 1000 μm. (B) Experimental replicate of *S. typhimurium* colonization dynamics in a spheroid. (C) Total sfGFP of bacteria (green) and spheroid size (purple) over time. Day 0 values are after inoculation and washing of bacteria. Error bars indicate ± s.e. averaged over three measurements. arbitrary unit. (D, E) Bacteria (purple) detected in the necrotic region in tumor spheroid via gram staining. Scale bar, 100 μm (D) and 10 μm (E).

**Fig. S3.**
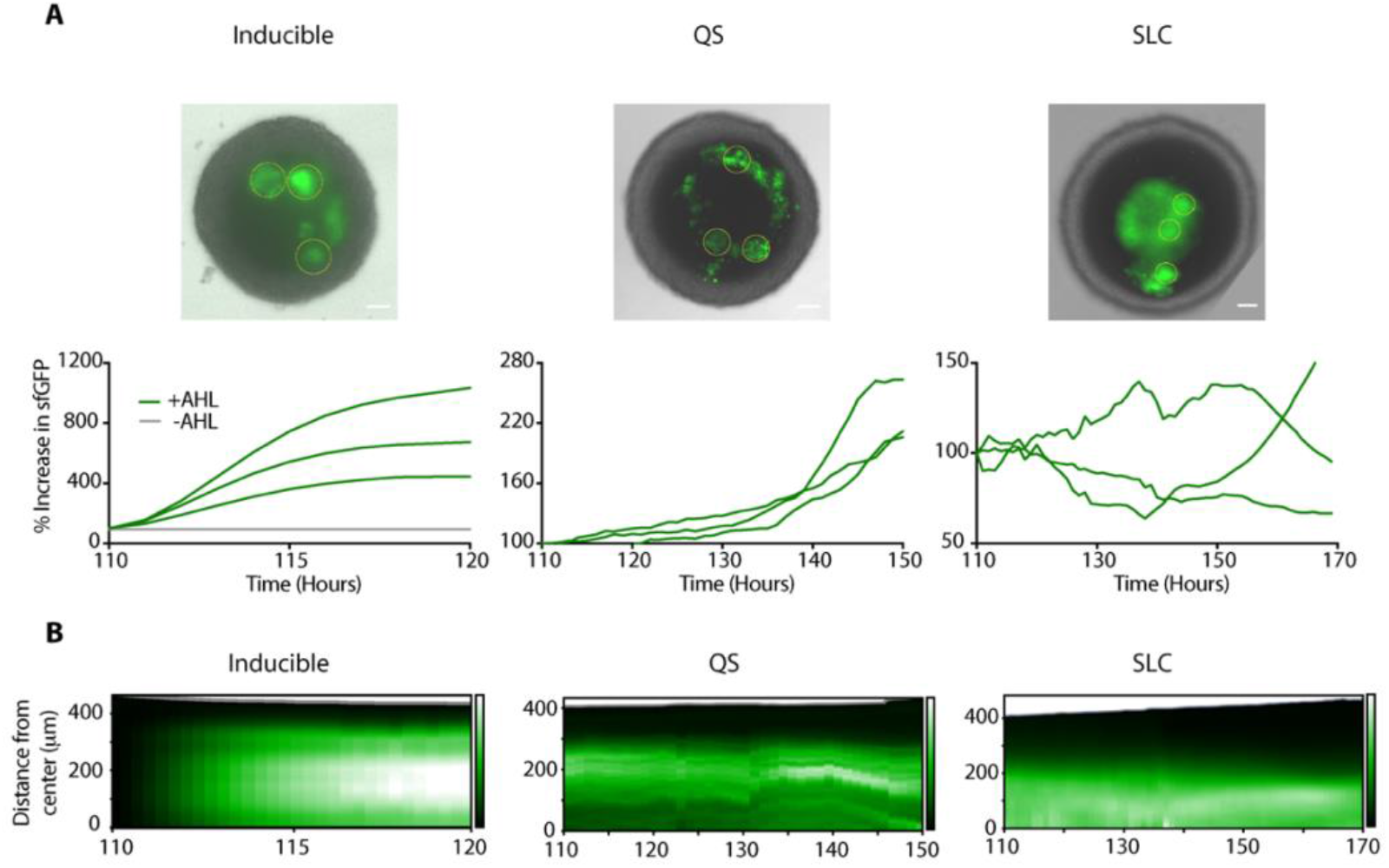
Automated image analysis of synthetic gene circuit dynamics in a spheroid. (A) (top) Circular region of interest (ROI’s) used to compute sfGFP fluorescence trajectories in spheroids. Scale bar, 100 μm. (bottom) Quantification of sfGFP signals. Signals are from 3 ROIs within a spheroid (see Methods). (B) Experimental replicates of synthetic gene circuits in spheroids.

**Fig. S4.**
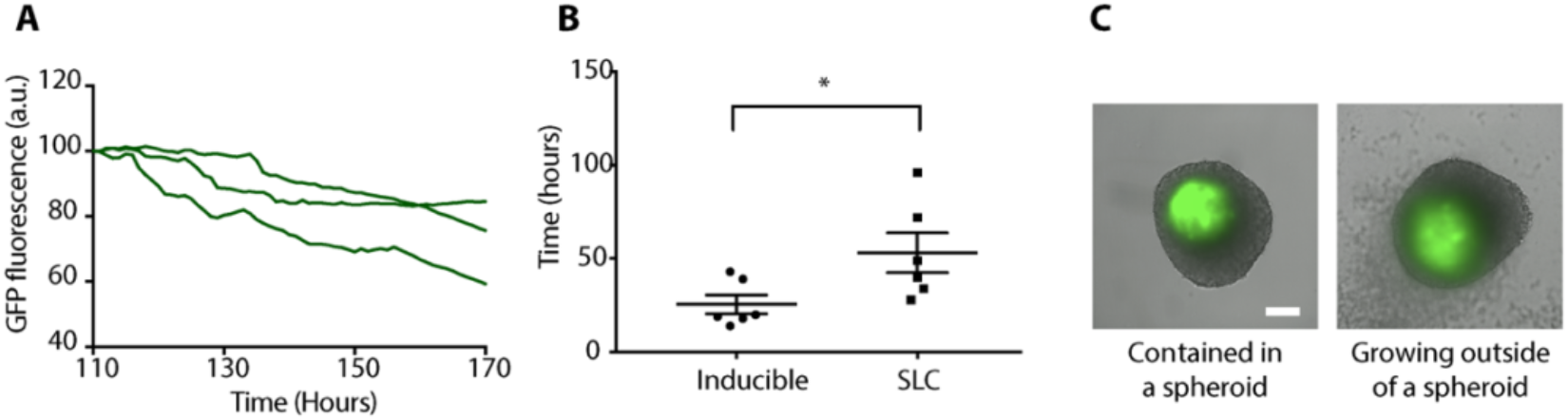
Biocontainment of SLC bacteria within tumor spheroids. (A) Total sfGFP of SLC bacteria over time. Signals are from three separate spheroids. (B) Time for bacteria to escape containment and grow outside of tumor spheroid (*P=0.0401, student’s t-test, error bars show s.e.). The time was determined as first frame to show detectable bacterial cells outside of spheroid boundary under transmitted light. (C) Typical images of sfGFP-expressing bacteria contained in tumor spheroids, or growing outside after removal of gentamicin from the media, respectively. Scale bar, 200 μm.

**Fig. S5.**
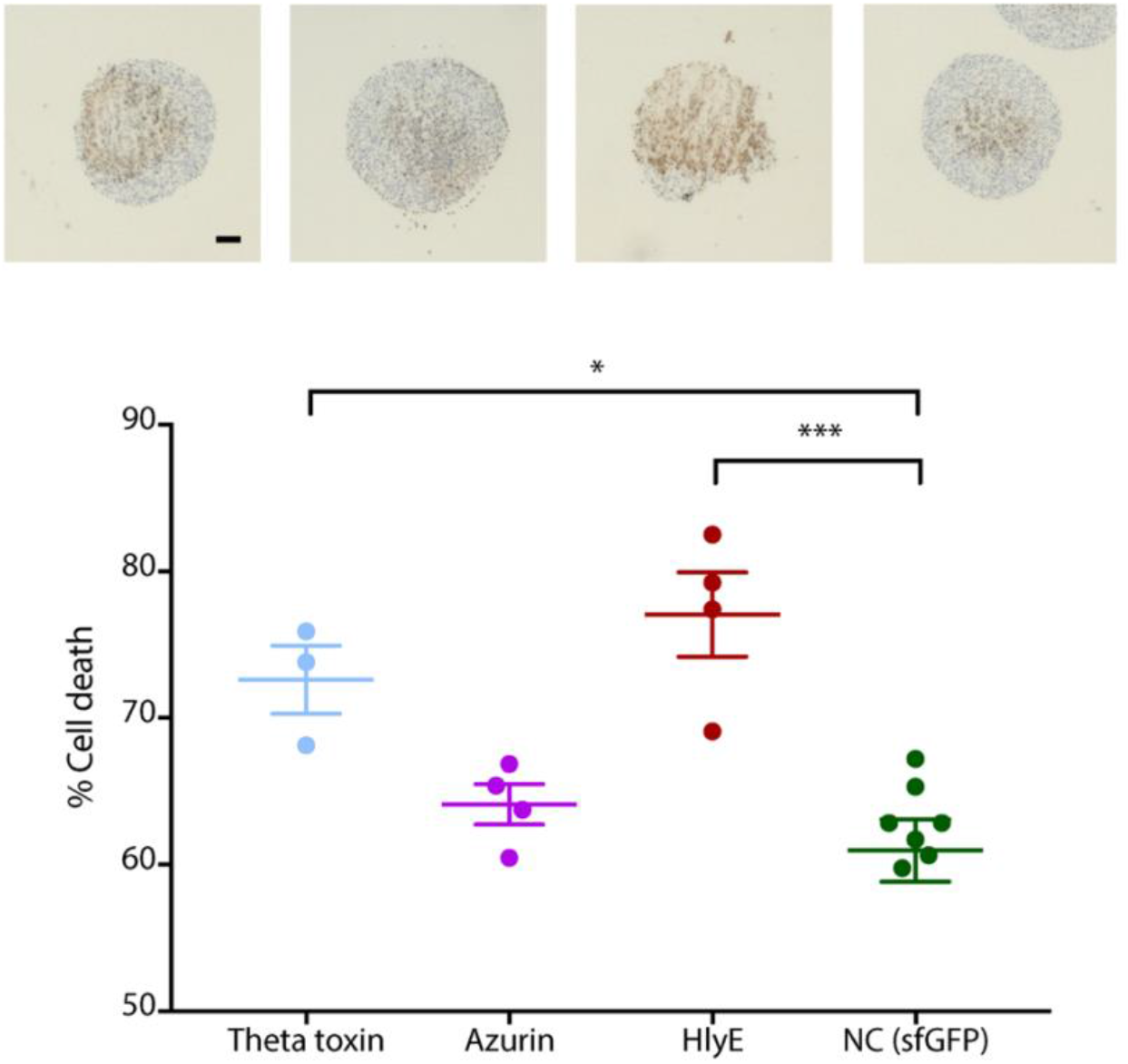
Histological analysis of tumor spheroid sections 10 days post bacteria inoculation. (top) Representative images of TUNEL staining for tissue sections. Scale bar, 100 μm. (bottom) Percentage cell death of histological tumor sections treated with bacteria expressing inducible therapeutics with 10 nM AHL 10 days post administration. TUNEL staining for tissue sections indicating apoptotic regions of tumors were used to obtain a measure of live and dead regions (*P=0.0112, ***P=0.0003, one-way ANOVA with Bonferroni post-test, error bars indicate ± s.e. averaged over > three measurements).

**Fig. S6.**
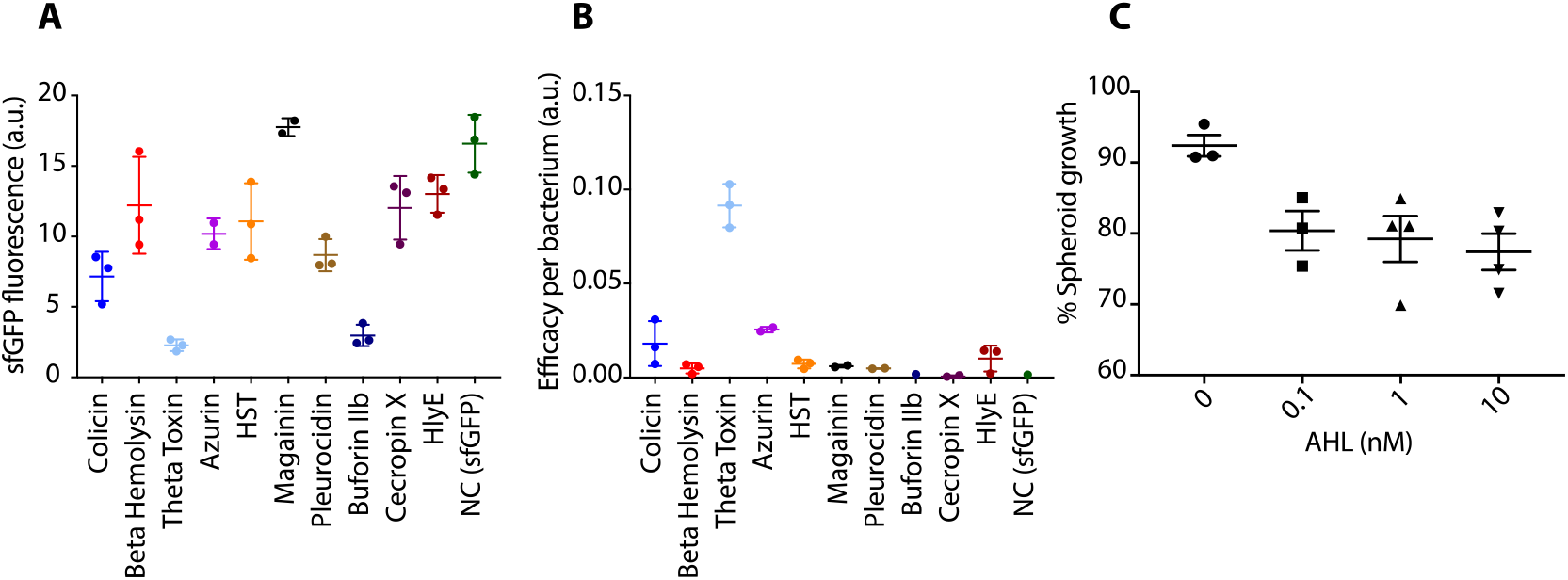
Effect of therapeutic expression on bacterial growth in tumor spheroids. (A) Bacteria growth in tumor spheroid 10 days post colonization measured by sfGFP fluorescence. Therapeutic expression was induced 4 days post bacteria inoculation with 10 nM AHL. (B) Efficacy of therapeutics normalized to bacteria optical density (OD). Error bars indicate ± s.e. averaged over 2-3 measurements. (C) Tumor spheroid growth treated with *S. typhymurium* expressing theta-toxin carrying inducible circuit with gradient of AHL induction (0, 0.1, 1, 10nM) 4 days post bacteria inoculation. Bacteria growth was measured by constitutively expressed sfGFP signal. The values are normalized by control measurement (bacteria expressing sfGFP) post 8 days of bacteria administration.

**Fig. S7.**
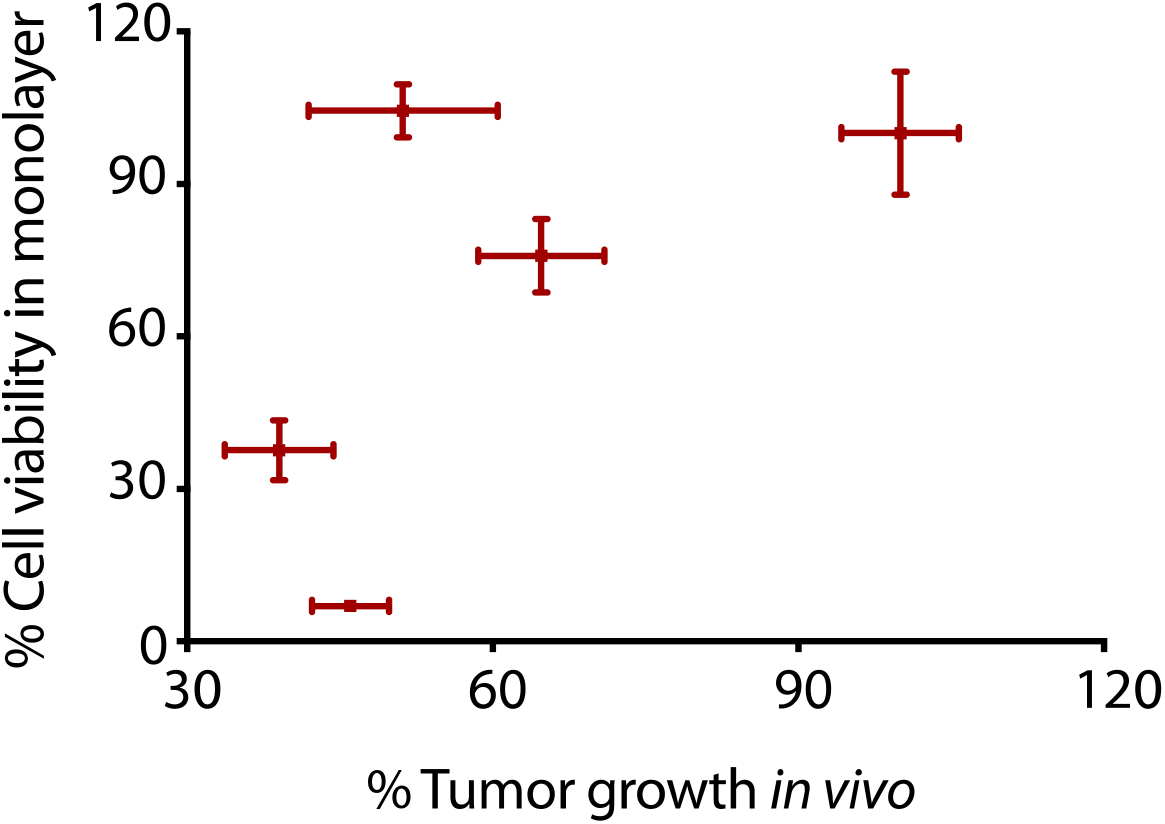
Comparison of efficacy between tumor growth in monolayer and *in vivo* model. The plot shows analysis of correlation (Linear regression, R^2^=0.38) treated with *S. typhimurium* carrying an inducible gene circuit expressing beta hemolysin, theta-toxin, azurin, hemolysin E, and sfGFP (control).

**Fig. S8.**
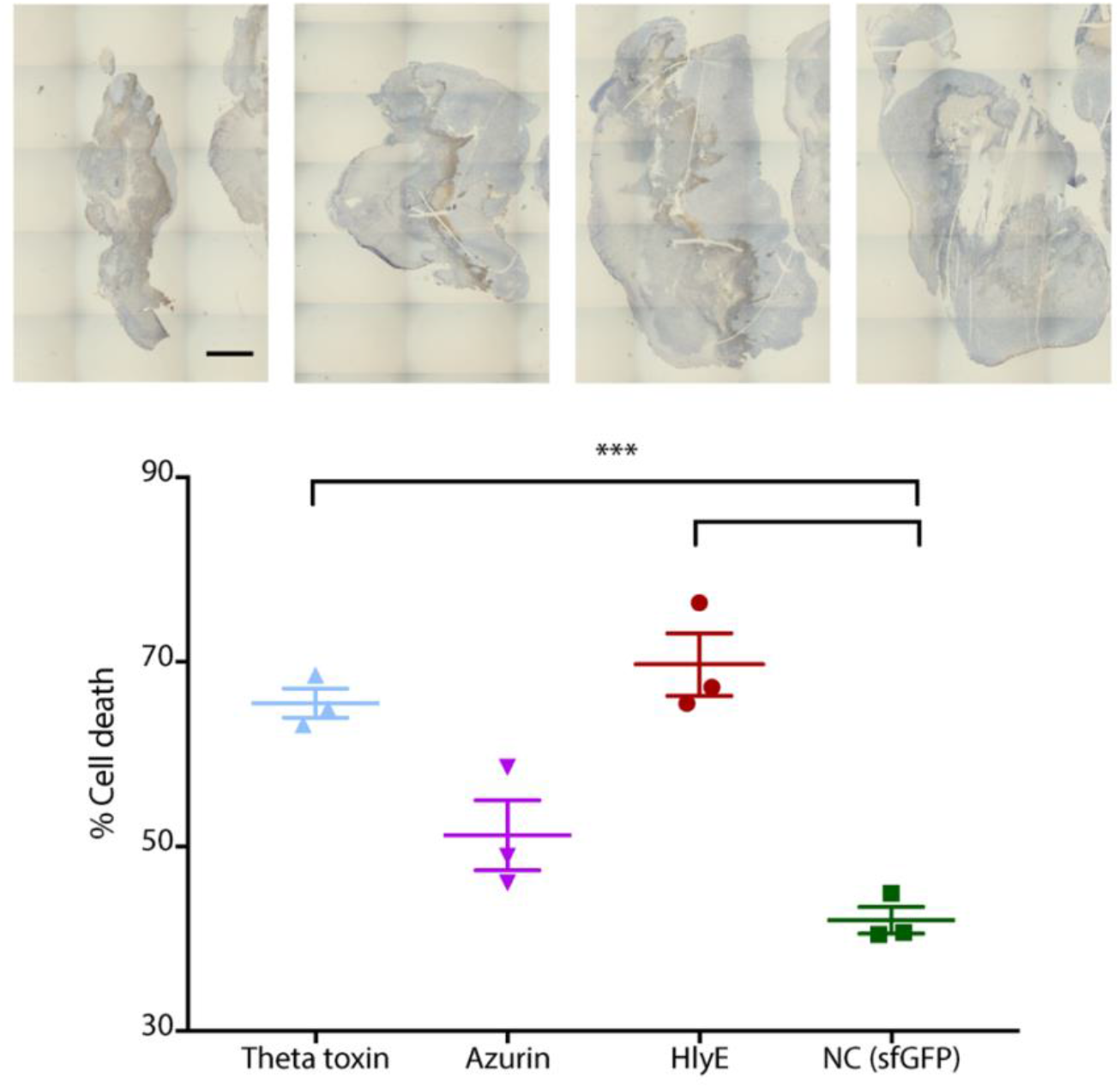
Histological analysis of tumor sections taken from mice 14 days post initial treatment. (top) representative images of TUNEL staining for tissue sections. Scale bar, 1 mm. (bottom) Percentage cell death of histological tumor sections treated with bacteria expressing inducible therapeutics 14 days post administration. TUNEL staining for tissue sections indicating apoptotic regions of tumors were used to obtain a measure of live and dead regions (***P=0.0009 theta toxin, ***P=0.0003 HlyE, one-way ANOVA with Bonferroni post-test, error bars indicate ± s.e. averaged over three measurements).

**Fig. S9.**
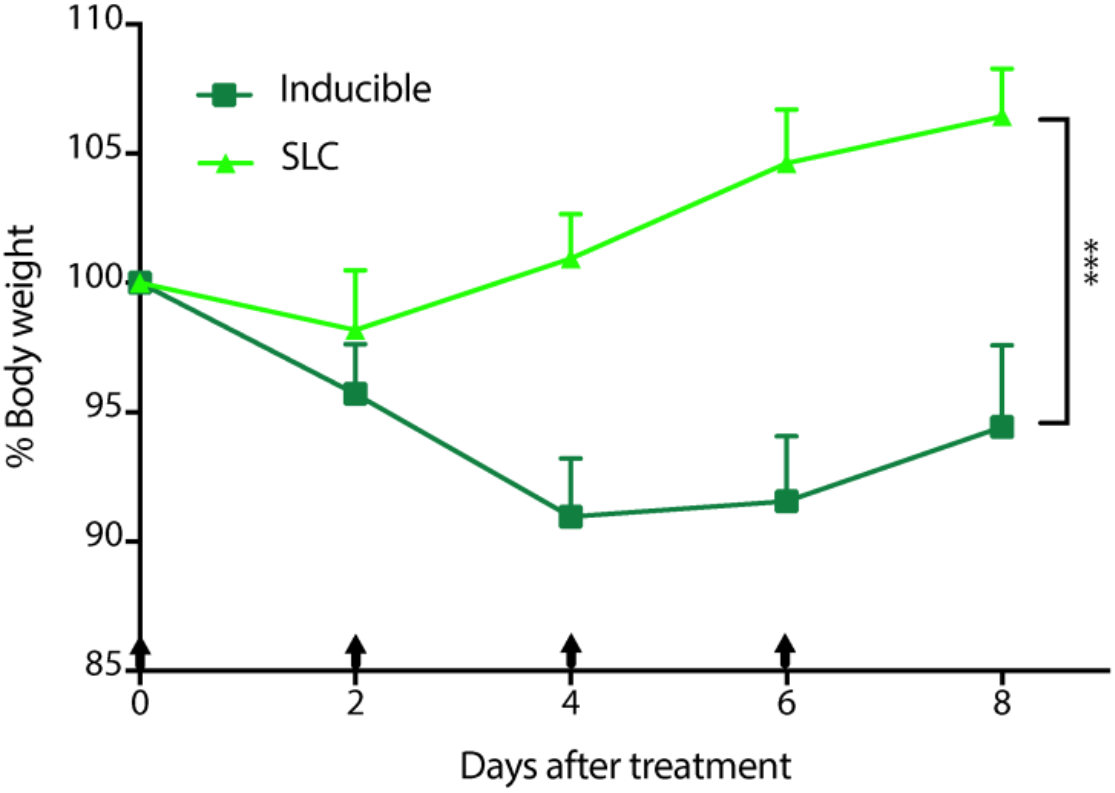
Percent change in body weight over time for the mice with subcutaneous tumor injected with *S. typhimurium* carrying inducible gene circuit and synchronized lysis circuit expressing sfGFP. Black arrows indicate bacteria injections (***P=0.0008, two-way ANOVA with Bonferroni post-test, n=5 mice, error bars show s.e.).

**Fig. S10.**
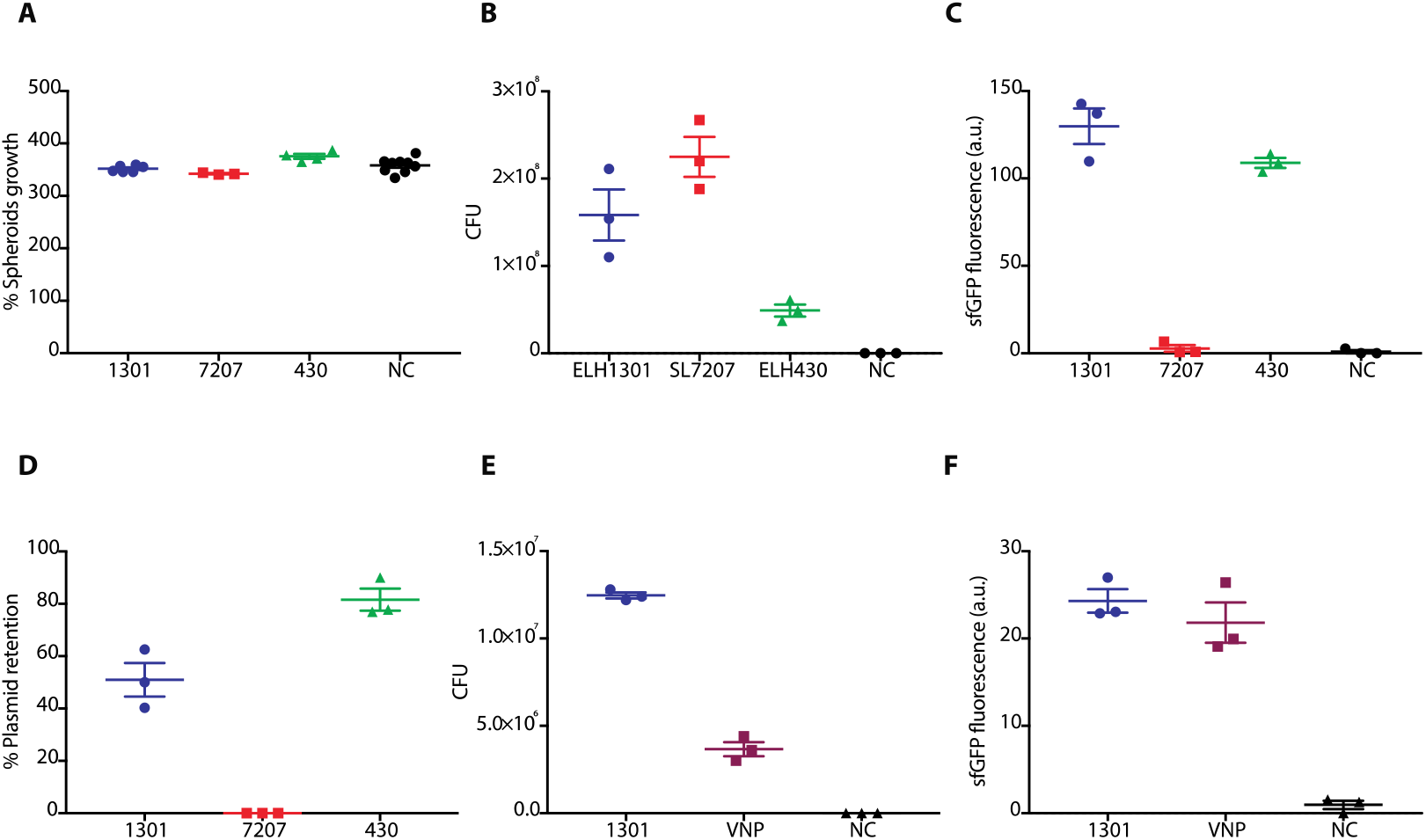
Comparison of *S. typhymurium* strains in tumor spheroids. (A) Tumor spheroid growth, (B) CFU of *S. typhymurium* strains (ELH1301, SL7207, and ELH430) recovered from dissociated spheroid and (C) sfGFP fluorescence of *S. typhymurium* strains in tumor spheroids, and (D) % plasmid retention recovered from dissociated tumor spheroid, 9 days post bacteria inoculation. Plasmid retention was calculated by colony counts on either plasmid-selective (kanamycin) or non–plasmid-selective plates. Error bars indicate ± s.e. averaged over three measurements. NC, negative control. (E, F) CFU of *S. typhymurium* strains (ELH1301 and VNP20009) recovered from tumor spheroids (E) and sfGFP fluorescence from *S. typhymurium* strains in tumor spheroids (F), 6 days post bacteria inoculation. Error bars indicate ± s.e. averaged over three measurements. NC, negative control.

**Fig. S11.**
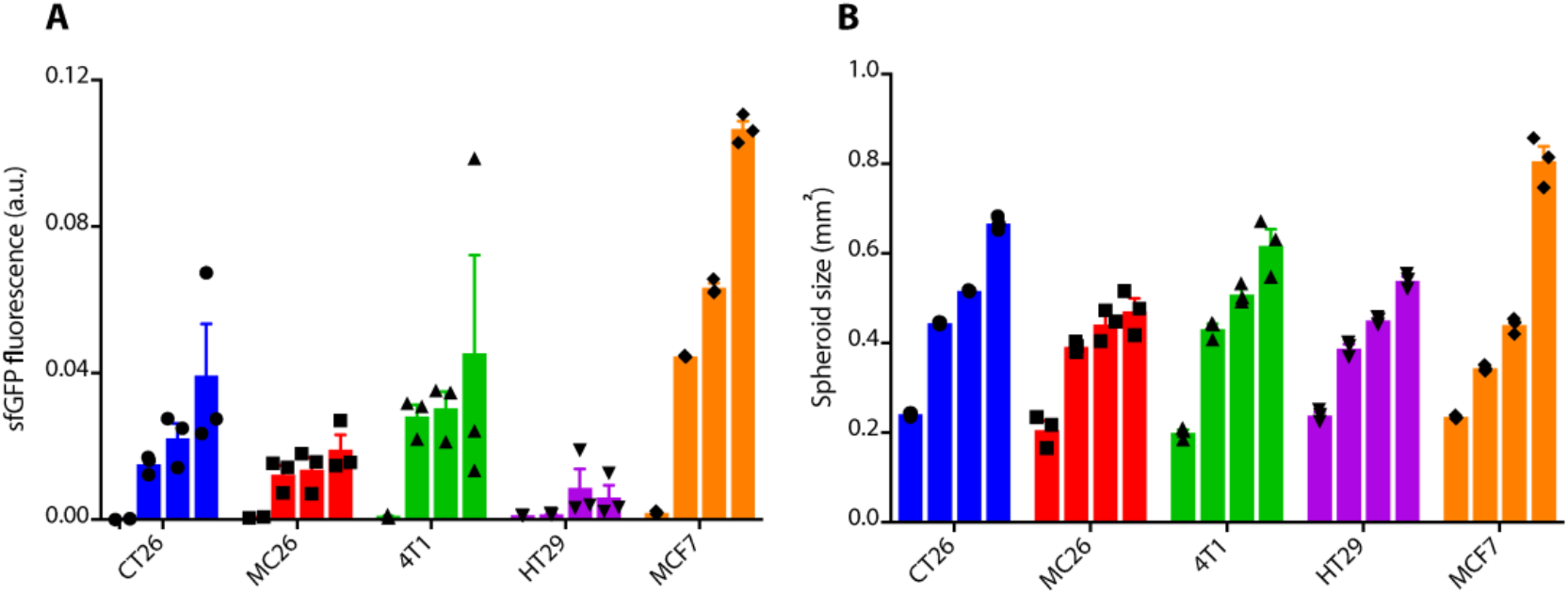
*S. typhimurium* growth in tumor spheroids derived from multiple cell types. (A) Bacteria growth over time in tumor spheroids as measured by sfGFP expression. (B) Tumor spheroid growth over time after colonization by bacteria. Bars indicate days 0, 2, 4, 6 post inoculation of bacteria. Error bars indicate ± s.e. averaged over three measurements.

**Fig. S12.**
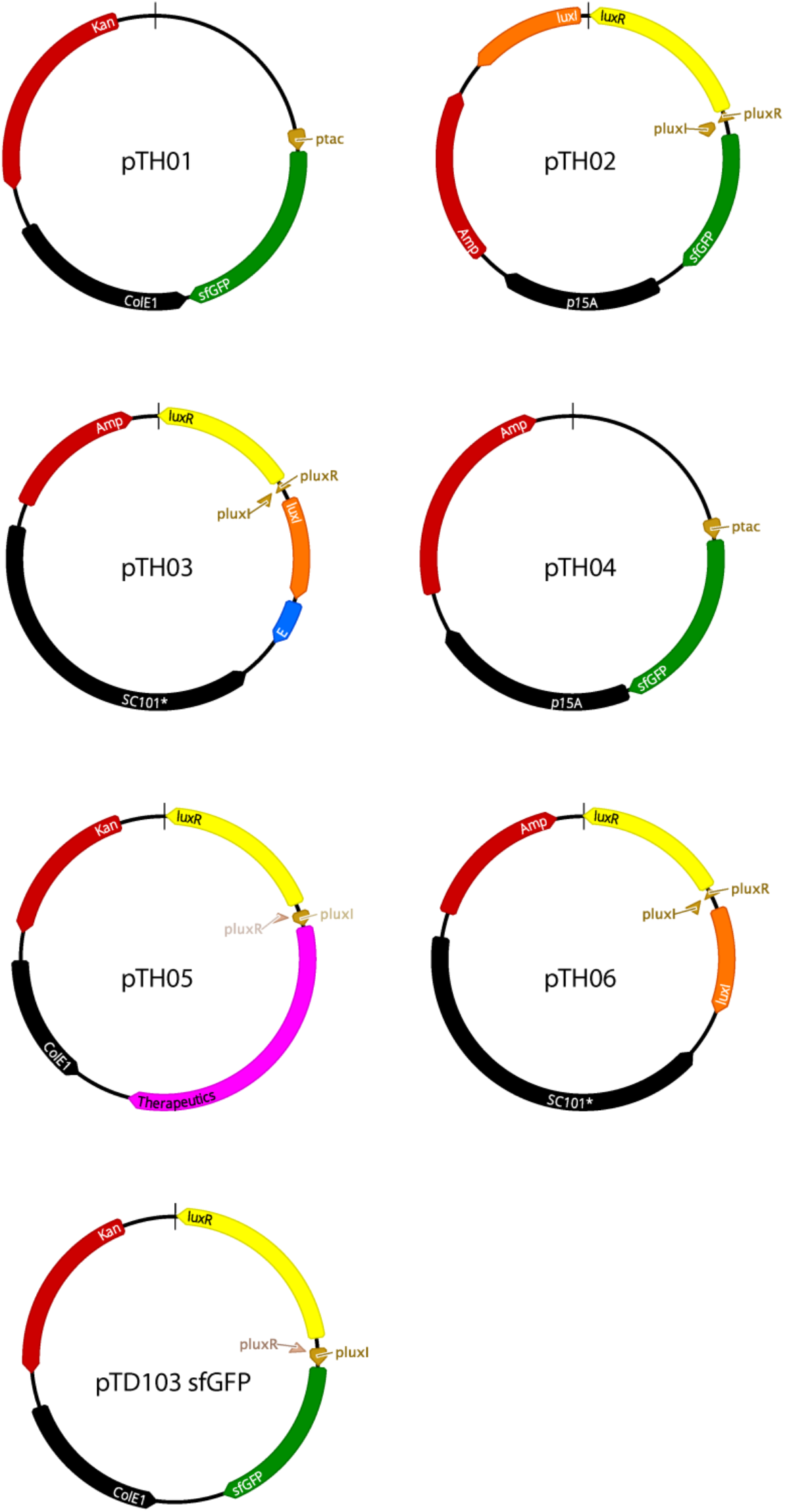
Map of main plasmids used in this study.

## Supplementary Tables

**Table S1.** A list of antimicrobial agents tested in this study.

| Categories | Antimicrobials | Tested (working) concentrations | Properties | References |
| --- | --- | --- | --- | --- |
| Antibiotics | Gentamicin | 0.5-50 µg/mL (5 µg/mL) | Reduced permeability to mammalian cell membrane, commonly used for antibiotic protection assay | Elsinghorst. 1994 |
|  | Erythromycin | 25-500 µg/mL (250 µg/mL) | High molecular weight, low accumulation in mammalian cellular cytosol | Kaneko et al., 2016<br>Carryn et al., 2003 |
|  | Tetracycline | 1-100 µg/mL (10 µg/mL) | High molecular weight, low accumulation in mammalian cellular cytosol | Kaneko et al., 2016<br>Carryn et al., 2003 |
| Antimicrobial peptides | Pyrrhocorcin | 0.1-60 µM (8-32 µM) | Reduced permeability to mammalian cell membrane due to proline-rich sequence | Hansen et al., 2012 |
|  | Apidaecin Ib | 0.1-60 µM (0.5-64 µM) | Reduced permeability to mammalian cell membrane due to proline-rich sequence | Hansen et al., 2012 |
|  | Drosocin | 0.1-60 µM (2-32 µM) | Reduced permeability to mammalian cell membrane due to proline-rich sequence | Hansen et al., 2012 |
| Metals | Silver Nanoparticle (100 nm diameter) | 1-50 µg/mL (3.12 µg/mL) | Reduced permeability to mammalian cell membrane due to large size | Albanese et al., 2013<br>Zarei et al., 2013 |

**Table S2.**
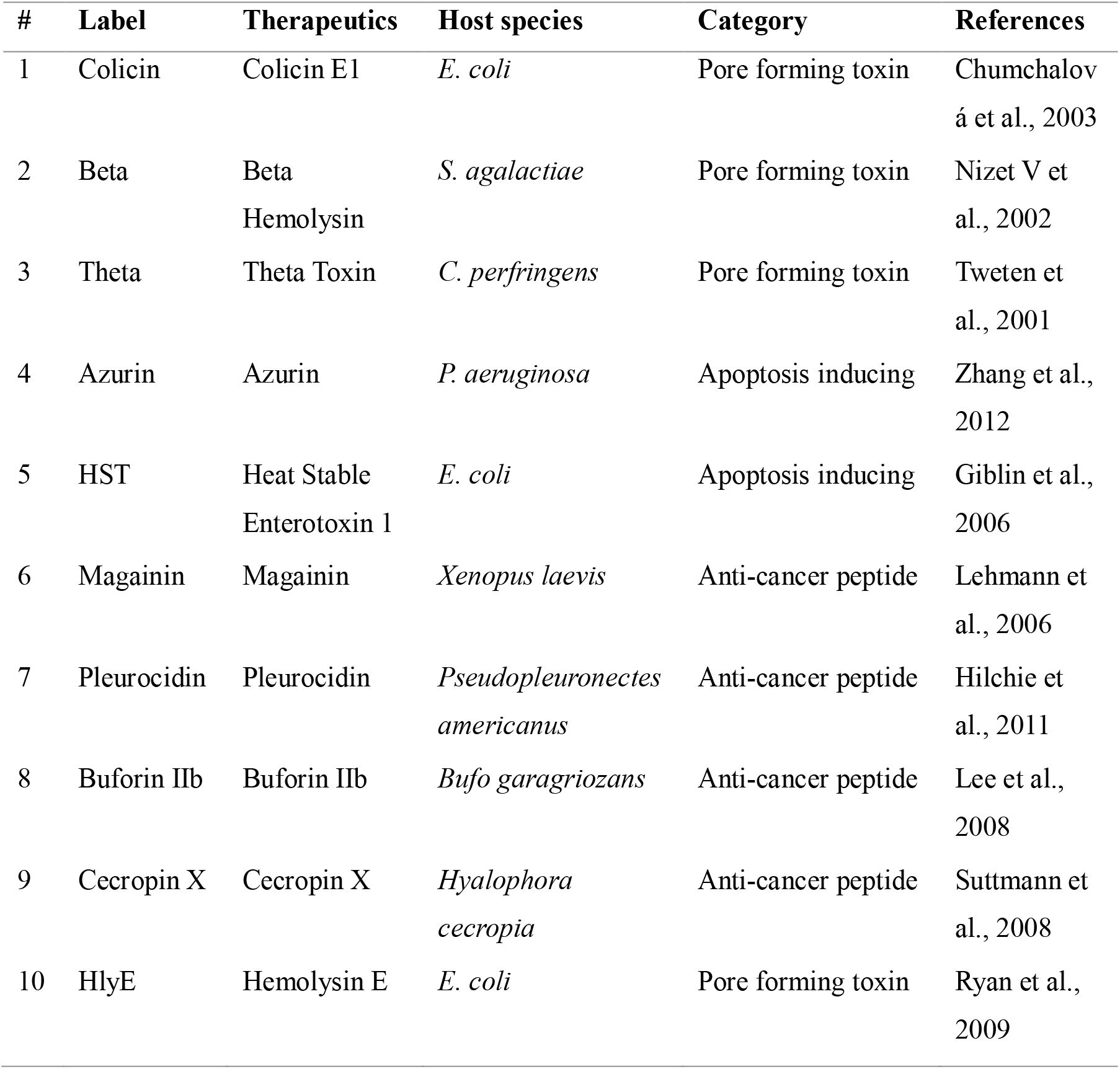
A list of therapeutics tested in this study.

**Table S3.** Co-culture protocols used in this study.

| **Bacteria species** | **Inoculation CFU** | **Incubation time**<br>**(hours)** | **Gentamicin**<br>**concentration (µg/mL)** |
| --- | --- | --- | --- |
| *S. typhimurium* | 10<sup>6</sup> | 2 | 2.5 |
| *L. monocytogenes* | 3 x 10<sup>7</sup> | 1 | 10 |
| *P. mirabilis* | 5 x 10<sup>5</sup> | 4 | 4 |
| *E. coli* | 3 x 10<sup>6</sup> | 6 | 1.5 |

**Fig. S4.**
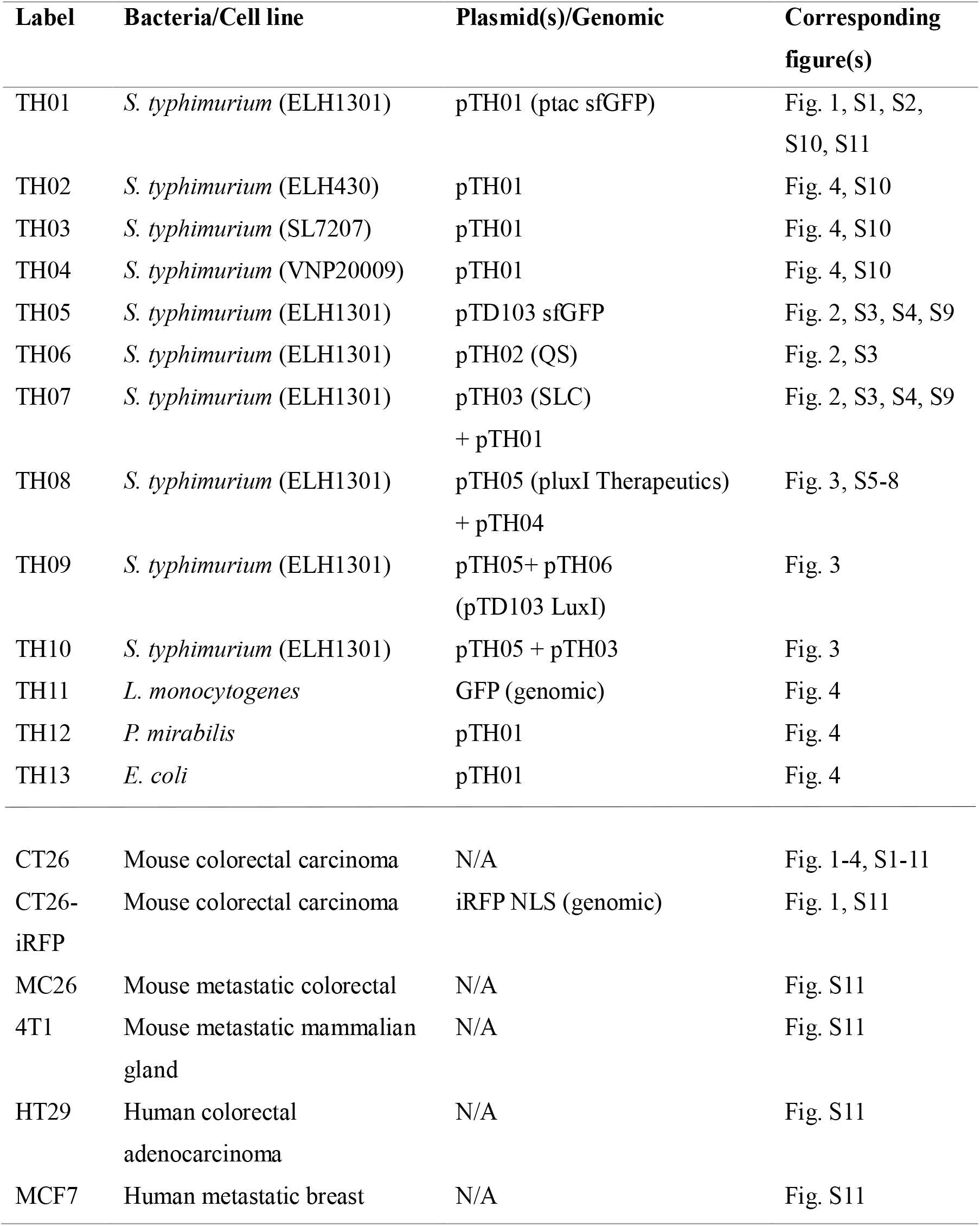
A list of bacteria and mammalian cells used in this study with respective plasmids/genomic changes.

## Supplementary Movie Legends

**Movie S1. Tumor spheroid formation.**

Time-lapse fluorescence microscopy of tumor spheroid formation. Images were taken every 30 minutes in low adhesion 96-well plate at 4X magnification.

**Movie S2. Dynamics of bacteria colonization of spheroids.**

Time-lapse fluorescence microscopy of *S. typhymurium* colonization of tumor spheroid. ELH1301 was inoculated to a 4-day old tumor spheroid, washed and incubated with gentamicin. Images were taken every 1 hour at 4X magnification.

**Movie S3. Inducible bacteria gene circuits in spheroids.**

Time-lapse fluorescence microscopy of inducible circuit in ELH1301 within tumor spheroid. ELH1301 carrying inducible circuit was induced by adding 10nM AHL to the media 4 days post spheroid colonization. Images were taken every 20 minutes at 4X magnification.

**Movie S4. Quorum-sensing bacteria gene circuit in spheroids.**

Time-lapse fluorescence microscopy of quorum-sensing circuit in ELH1301 within tumor spheroid. The strain was inoculated to 4-day old tumor spheroid and images were taken every 1 hour at 4X magnification.

**Movie S5. Synchronized lysis bacteria gene circuit in spheroids.**

Time-lapse fluorescence microscopy of Synchronized Lysis Circuit (SLC) in ELH1301 within tumor spheroid. The strain was inoculated to 4-day old tumor spheroid and images were taken every 1 hour at 10X magnification.

